# Primate Prosocial Behaviour as Socially Gated Action Selection

**DOI:** 10.64898/2026.07.14.737163

**Authors:** Christoph D. Dahl

## Abstract

Research on prosocial behaviour in nonhuman primates faces two related but distinct problems. First, empirical findings are heterogeneous: studies report instrumental helping, targeted tool transfer, food provisioning, consolation, and collaborative action, whereas others report little or no concern for partner welfare in low-cost prosocial-choice paradigms. These differences may reflect paradigm-specific variation in recipient need, action cost, actor benefit, joint payoff, solicitation, relationship context, competition, affordance clarity, and response bias rather than disagreement about one mechanism. Second, the interpretive vocabulary is underconstrained. Terms such as altruism, empathy, prosocial motivation, helping, and collaboration are often defined from behavioural outcomes and then treated as if they identified distinct latent mechanisms. This manuscript proposes a computational reframing of the problem. Prosocial behaviour is formalised as socially gated action selection. In the model, instrumental helping, low-cost prosocial choice, costly recipient benefit, strict altruism-like outcomes, need-sensitive empathy-like responding, and coordinated joint action are task and outcome profiles generated by different configurations of independently specified variables. These include need detection, goal inference, affordance recognition, action cost, actor benefit, relationship and tolerance effects, competition, request context, social feedback, joint payoff, action selection, and motor or apparatus bias. An illustrative simulation shows how low-cost instrumental helping, cost- or risk-suppressed recipient benefit, false-positive prosocial choice, and reliable-partner collaboration can arise from the same architecture. The framework identifies which task variables must be measured or manipulated before stronger claims about other-regarding motivation, empathy-like responding, or collaboration are justified.

## Introduction

Prosocial behaviour in nonhuman primates is behaviourally observable but mechanistically underdetermined. An actor may produce an action that improves another individual’s state, yet similar recipient-benefiting outcomes can arise from different processes, including detection of another’s need, recognition of an effective affordance, low action cost, reduced competition, relationship tolerance, solicitation, direct actor benefit, joint payoff, motor bias, or other-regarding valuation. This creates a first problem: empirical findings across instrumental helping, food transfer, consolation, prosocial choice, and collaboration need not converge, because these paradigms differ in the task variables that make recipient benefit possible or costly. A second problem concerns interpretation. Terms such as altruism, empathy, consolation, sharing, helping, and collaboration are useful as descriptive labels, but they become problematic when behavioural outcomes are treated as direct evidence for internal mechanisms. An animal that acts in a way that benefits another individual has produced a recipient-benefiting outcome. It does not follow, without further evidence, that the action was generated by altruistic motivation, empathic concern, or a representation of another’s welfare. This distinction is especially important in primate prosociality, where similar behavioural outcomes can reflect selfish, mutualistic, relationship-mediated, or other-regarding processes (de Waal and Suchak, 2010).

The empirical literature illustrates the problem. Instrumental-helping studies have shown that young chimpanzees can assist a human experimenter in transparent goal contexts (Warneken and Tomasello, 2006). Tool-transfer experiments have further shown that chimpanzees can provide a conspecific with a useful object, especially when the recipient’s problem is visible and when the recipient requests assistance (Yamamoto et al., 2009, 2012). At the same time, prosocial-choice studies have produced inconsistent results. Some studies concluded that chimpanzees failed to deliver low-cost benefits to familiar conspecifics (Silk et al., 2005; Jensen et al., 2006), whereas other designs elicited more prosocial choices or showed that social and task structure strongly affect the outcome (Horner et al., 2011; House et al., 2014; Mendonça et al., 2018; Sikorska et al., 2026). These task effects are not merely technical. Mendonça et al. (Mendonça et al., 2018) are especially relevant because extended touch-screen-guided testing allowed the prosocial-choice task to be sampled over more trials than in earlier procedures (Horner et al., 2011), revealing higher rates of prosocial choice. At the same time, dyadic variation indicated that prosocial choice should not be treated as a unitary trait, but as a context-sensitive outcome shaped by relationship, rearing history, and partner-specific valuation. Conversely, Sikorska et al. (Sikorska et al., 2026) show that physical token arrangement can alter apparent prosocial responding, with fixed token positions producing choices shaped by reaching convenience and lateralisation. Cooperative breeders such as common marmosets provide a different contrast, because they appear more likely than chimpanzees to provision others in some food-giving paradigms (Burkart et al., 2007), consistent with broader accounts that combine natural history and experimental evidence in explaining primate prosociality (Jaeggi et al., 2010). The field is therefore not simply divided between positive and negative evidence. Rather, prosocial outcomes appear to depend on information access, solicitation, affordance clarity, resource context, relationship, social tolerance, partner identity, and response bias.

The present framework starts from a conservative premise. The unit of explanation should not be altruism, empathy, or prosociality as an assumed faculty. The unit of explanation should be an action-selection process in which the actor integrates self-benefit, other-benefit, joint payoff, cost, relationship value, competition, request signals, and task affordances. The traditional labels are then treated as observer-level classifications of the resulting action and outcome. This allows one mechanism to account for several behavioural categories without assuming that these categories are themselves mechanisms. The central proposal is therefore:

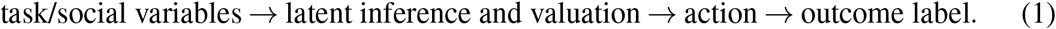

In compact form, the framework asks which conditions must be satisfied before a recipient-benefiting outcome can be interpreted psychologically: was recipient need detected; was an effective affordance available and recognised; what cost did the actor face; did competition or dominance risk suppress action; did relationship value or tolerance alter partner valuation; did request signals or prior social feedback change the salience of the recipient’s state; and did action selection favour the recipient-benefiting option over available alternatives? These questions correspond to variables that must be measured or manipulated independently of the observed outcome. Recipient-need detection can be constrained by whether the recipient’s problem is visible or hidden. Affordance recognition can be constrained by whether the helping action is available, familiar, or perceptually clear. Cost and competition can be constrained by effort, delay, food loss, dominance risk, or active food competition. Relationship and tolerance effects can be constrained by partner identity, affiliation, kinship, or rearing history. Request and feedback effects can be constrained by solicitation, reaching, begging, or previous social interaction. Action-selection effects can be constrained by the available alternatives and by controls for side, hand, location, motor convenience, and apparatus bias. This causal direction is essential. A circular model would infer a helping gate from the fact that helping occurred. A non-circular model instead defines the gates from independently measured or experimentally manipulated variables and then asks whether those variables predict recipient-benefiting action.

### Scope and aim of the framework

This article develops a theoretical and generative modelling framework for prosocial behaviour in nonhuman primates. It is not intended as a catalogue of all reports of helping, sharing, consolation, provisioning, prosocial choice, or cooperation, nor does it attempt to provide an empirical model of any single species. The aim is methodological: to show how behavioural categories and interpretive labels such as instrumental helping, prosocial choice, empathy-like responding, costly recipient benefit, altruism-like outcome, and collaboration can be decomposed into independently measurable task and valuation components. Representative paradigms are used to identify recurrent inferential problems in the field: whether the actor detected recipient need, whether the actor understood the relevant goal, whether an effective affordance was recognised, whether cost or competition suppressed action, and whether response bias could mimic prosocial choice. This makes it possible to ask which experimental conditions distinguish other-state-sensitive action from low-cost responding, task engagement, competition, relationship effects, or apparatus bias. The intended use is therefore not a general theory of animal prosociality, but an inferential framework for interpreting primate instrumental helping, tool transfer, token choice, food sharing, provisioning, post-conflict affiliation, prosocial choice, and cooperation. A fuller mapping of the empirical literatures onto the proposed gating and valuation terms is provided in the Supplementary Material, Table S1.

### From labels to action selection

The proposed model does not deny that terms such as helping, prosociality, altruism, empathy, or collaboration can be useful. It denies that these terms should be used as explanatory primitives. They are better treated as different regions of a common action-selection space. Table 1 summarises this replacement. In the present manuscript, helping is used as the broadest recipient-benefit outcome term: an action is helping when it improves the recipient’s state. Prosocial choice is treated as a narrower low-cost subset of helping, whereas costly other-benefit, strict altruism-like outcome, empathy-like responding, and collaboration impose additional or distinct diagnostic requirements. For the purpose of the table, *a* denotes the selected action, Δ*X*_*j*_(*a*) the realised change in the recipient’s state, Δ*X*_*i*_(*a*) the realised change in the actor’s state, *C*_*i*_(*a*) the action cost paid by the actor, and *ε* a small threshold below which the action is treated as low-cost. *P*_*i*_(*a*_*help*_ | *z*_*t*_) denotes the actor’s probability of selecting the helping action in task context *z*_*t*_, and 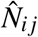denotes inferred recipient need. *X*_*i j*_ denotes joint payoff, and *a*_*i*_ and *a* _*j*_ denote actor and partner actions. The benefit of this formulation is that the same architecture can generate apparently different behaviours. Helping occurs when an action improves another individual’s state. Prosociality occurs when that benefit is low-cost. Costly other-benefit occurs when that benefit is accompanied by actor cost. Empathy-like responding occurs when action probability changes with inferred need. Collaboration occurs when the action value depends on predicted joint action. Strict altruism-like outcomes form a narrower subset in which the recipient benefits while the actor’s net outcome is negative. The mechanism is not a separate altruism module, empathy module, or collaboration module; it is socially modulated action selection.

**Table 1:**
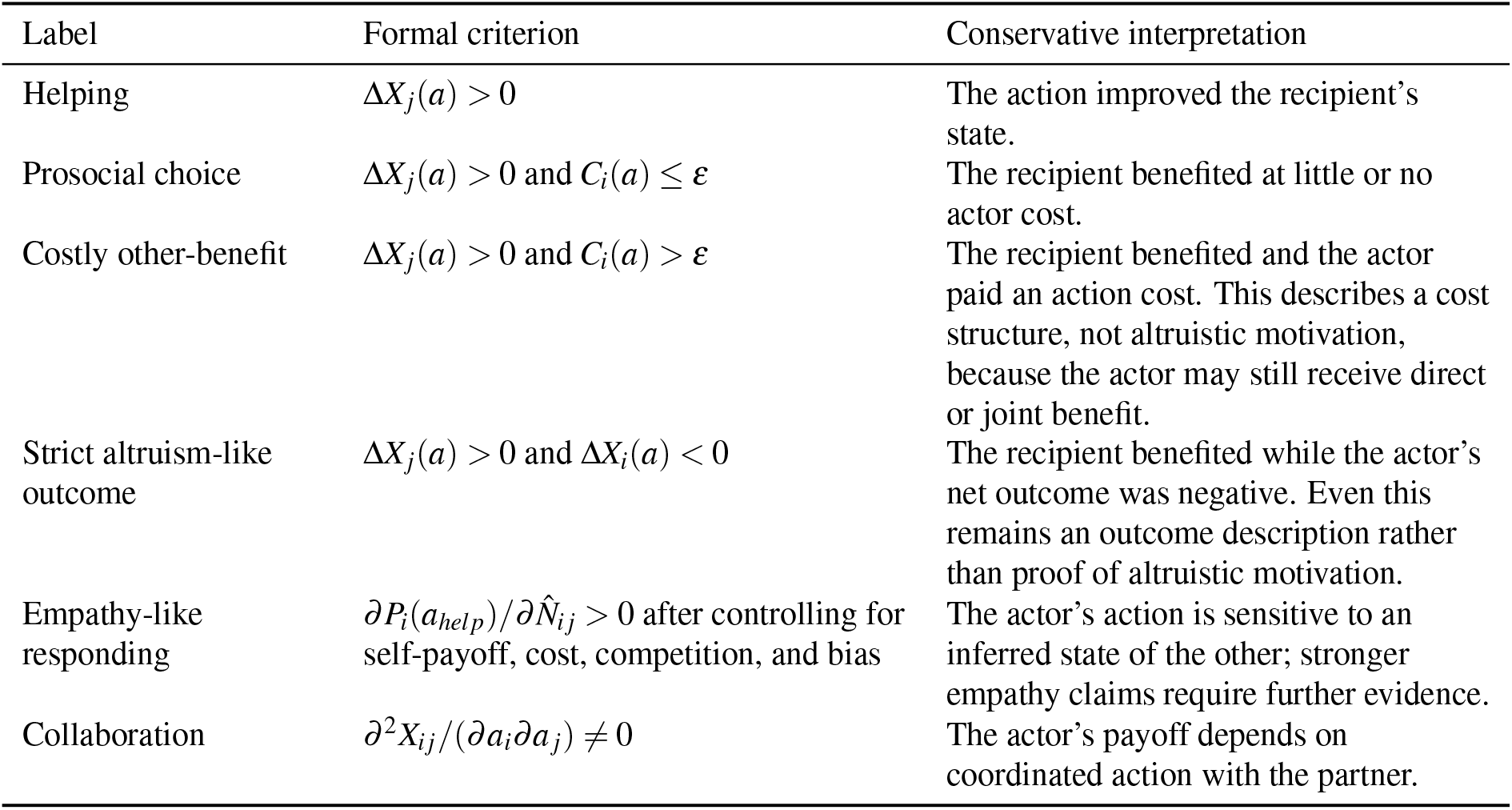
Traditional labels recast as outcome profiles of a common action-selection architecture.

## Materials and Methods

### Model overview

Let actor *i* face a possible recipient *j* in trial *t*. The actor can choose an action *a* from a finite action set *A*_*i*_. The task-social context is represented as:

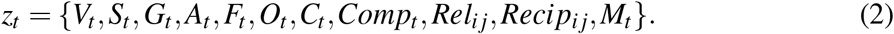

The variables are defined in Table 2. Importantly, these variables are not defined from whether helping occurred. They are independently measured, experimentally manipulated, or externally coded.

**Table 2:** Main variables in the social-gating model.

| Symbol | Meaning | Possible independent measurement or manipulation |
| --- | --- | --- |
| $V_t$ | Visibility of recipient need | Transparent versus occluded recipient problem. |
| $S_t$ | Solicitation or request intensity | Reaching, begging, vocalising, directed request, or experimentally prevented request. |
| $G_t$ | Goal transparency | Whether the recipient’s goal is obvious from behaviour and apparatus. |
| $A_t(a)$ | Affordance clarity for action $a$ | Actor’s prior solo success with the apparatus; physical action-outcome transparency. |
| $F_t$ | Task familiarity | Training history or number of prior trials. |
| $O_t$ | Occlusion or ambiguity | Visual obstruction, hidden apparatus state, or ambiguous recipient access. |
| $C_{i,t}(a)$ | Cost of action $a$ to actor | Distance, effort, risk, delay, food loss, or opportunity cost. |
| $Comp_t$ | Competition/resource conflict | Food context, dominance risk, co-feeding tolerance, or dyadic aggression. |
| $Rel_{ij}$ | Relationship or partner value | Grooming, proximity, kinship, tolerance, affiliation, dominance relation, partner identity, or alliance value (Cronin, 2012). |
| $Recip_{ij}$ | Reciprocal or interaction history | Prior received help, exchange history, repeated interaction, partner-specific history, or long-term dyadic association. |
| $M_t(a)$ | Motor/apparatus bias | Side bias, hand preference, motor lateralisation, token placement, and physical reaching convenience. |

### Need inference

The recipient has a true need state,

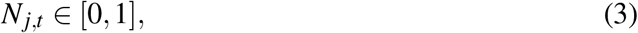

which is not directly available to the actor. The actor forms an inferred need estimate:

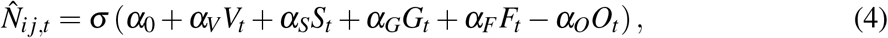

Where

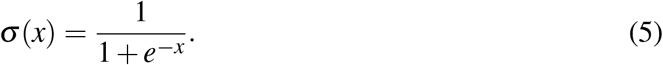

The coefficients *α*_*0*_, *α*_*V*_, *α*_*S*_, *α*_*G*_, *α*_*F*_, *α*_*O*_ are need-inference parameters. The intercept *α*_0_ sets baseline inferred need, whereas *α*_*V*_, *α*_*S*_, *α*_*G*_, *α*_*F*_, and *α*_*O*_ determine how strongly visibility, solicitation, goal transparency, task familiarity, and occlusion or ambiguity contribute to the actor’s inferred need estimate. Positive coefficients increase inferred need. The occlusion term is written with a negative sign because ambiguity is assumed to reduce need detection. The inferred-need estimate 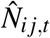captures the actor’s detection of recipient need. An actor may therefore fail to produce a recipient-benefiting action not because other-regarding valuation is absent, but because the relevant need state is not detected.

### Affordance recognition

Even if need is detected, the actor must identify an effective action. The actor’s perceived benefit to the recipient is:

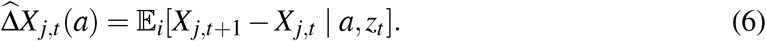

The operator *E*_*i*_[ ] denotes the actor’s subjective expectation. Thus, 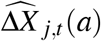 is not the realised benefit to the recipient, but the benefit that actor *i* expects action *a* to produce for recipient *j* in context *z*_*t*_. The probability that action *a* is represented as an effective helping affordance is:

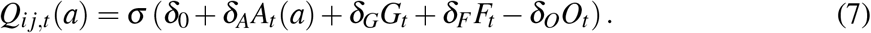

The coefficients δ_0_, δ_*A*_, δ_*G*_, δ_*F*_, δ_*O*_ are affordance-recognition parameters. The intercept δ_0_ sets baseline affordance recognition, whereas δ_*A*_, δ_*G*_, δ_*F*_, and δ_*O*_ determine how strongly affordance clarity, goal transparency, task familiarity, and occlusion or ambiguity influence whether the actor represents action *a* as effective. Need detection and affordance recognition are combined into a social gate:

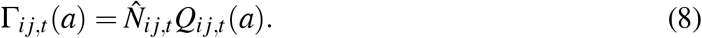

The gate is near zero if the actor does not detect the recipient’s need or does not recognise which action would help. This is the central anti-circular distinction: the gate is derived from visibility, request, goal transparency, familiarity, occlusion, and affordance clarity, not from the eventual helping outcome.

### Valuation and action selection

A central distinction is made between direct actor benefit, recipient benefit, and non-additive joint benefit. Direct actor benefit, 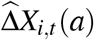, refers only to outcomes obtained by the actor. Recipient benefit, 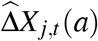, refers only to expected improvement in the recipient’s state. Joint benefit, 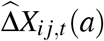, captures outcomes that depend on coordinated or mutually beneficial action. This separation is important because recipient benefit should not be smuggled into the actor’s self-interest term. A helping action can therefore benefit the recipient while providing no direct benefit to the actor, whereas a collaborative action can benefit both actor and recipient through a joint-payoff structure. For each action, the actor computes a utility:

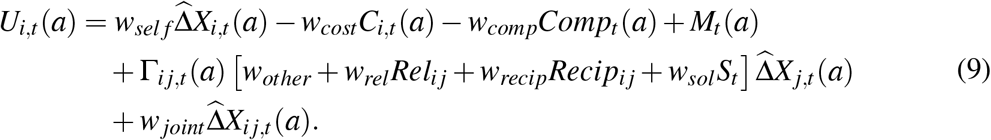

The weights determine how strongly each component contributes to action value. The term *w*_*sel f*_ weights direct benefit to the actor, *w*_*cost*_ weights action cost, and *w*_*comp*_ weights the suppressive effect of competition. The term *M*_*t*_(*a*) captures motor or apparatus bias in favour of action *a*. The parameter *w*_*other*_ weights baseline sensitivity to recipient outcome, whereas *w*_*rel*_, *w*_*recip*_, and *w*_*sol*_ modulate recipient benefit according to relationship value, prior reciprocal history, and current solicitation. Finally, *w*_*joint*_ weights non-additive joint benefit, as in coordinated or mutually beneficial action. In words, the first line asks whether the action benefits the actor, is costly, is suppressed by competition, or is favoured by motor or apparatus bias. The second line asks whether the action is socially gated: recipient benefit contributes to utility only to the extent that recipient need and an effective affordance are represented. Within this gated term, relationship, reciprocity, and solicitation can increase or decrease how strongly the recipient’s expected benefit affects the actor’s choice. The third line adds joint outcome value, which is relevant when coordinated action produces a benefit that is not reducible to separate actor and recipient payoffs. The term *w*_*other*_ should therefore be interpreted conservatively as other-outcome sensitivity. It is not, by itself, empathy or altruistic motivation.

Action selection follows a softmax rule:

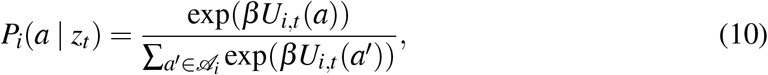

where β is choice consistency. Low β produces noisy choice; high β produces near-deterministic selection of the highest-valued action. The softmax rule therefore converts the utility of each available action into a choice probability, allowing the same valuation structure to generate variable behaviour rather than deterministic responding on every trial.

### Outcome and diagnostic classification

The previous subsections define how an action is valued and selected. The present subsection defines what may be said after the action has occurred. This step is classificatory rather than explanatory: realised outcomes are mapped onto descriptive labels, but those labels are not treated as mechanisms.

The world changes after action selection. The actual actor and recipient outcomes are:

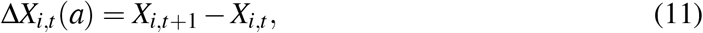

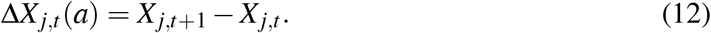

These realised outcomes are distinct from the expected outcomes 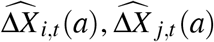, and 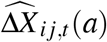 that entered the actor’s utility before the action was selected.

A recipient-benefiting outcome is present when:

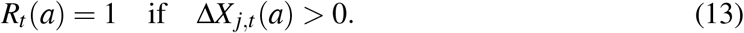

Depending on the task, such an outcome may correspond to instrumental helping, provisioning, food transfer, or another prosocial outcome. The label states what happened to the recipient. It does not by itself identify the actor’s motivation. The distinction between action cost and net actor outcome is important. The cost term *C*_*i,t*_(*a*) describes what the actor pays to perform the action. The net actor outcome Δ*X*_*i,t*_(*a*) describes the realised consequence for the actor after direct benefit, joint benefit, and cost have been combined. A costly recipient-benefiting action therefore does not necessarily imply a strict altruism-like outcome.

Prosocial choice is defined as low-cost recipient benefit:

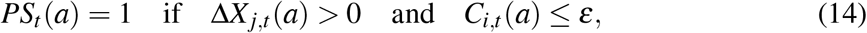

where *ε* is a small threshold for negligible action cost.

Costly recipient benefit is:

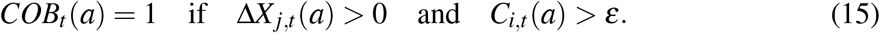

This label states that the recipient benefited while the actor paid an action cost. It should not be interpreted as altruism, because the actor may still obtain direct or joint benefit.

A stricter altruism-like outcome requires a negative net actor outcome:

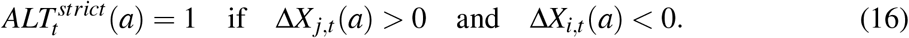

Empathy-like responding is not assigned merely because a recipient-benefiting action occurred. Unlike the preceding labels, it requires a counterfactual diagnostic: the probability of choosing the recipient-benefiting action must increase with inferred recipient need while alternative explanations are held constant:

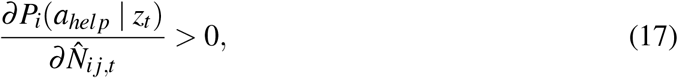

where *a*_*help*_ denotes the action that would benefit the recipient in the relevant task. This condition is deliberately weaker than a claim of empathy. It shows only that action probability changes with inferred recipient need, after direct self-benefit, cost, competition, relationship, and motor bias are held constant. It does not by itself establish empathic concern, emotional sharing, or a representation of the recipient’s welfare.

Collaboration requires non-additive joint payoff. If actor and partner choose actions *a*_*i*_ and *a* _*j*_, collaboration is present when:

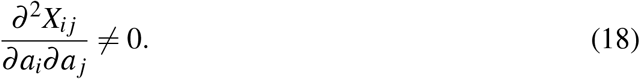

This expression states that the effect of one individual’s action on the joint outcome depends on the other individual’s action. The joint payoff is therefore not merely the sum of two independent individual payoffs; it contains an interaction between the partners’ actions. In discrete tasks this means that the joint outcome is not reducible to the separate actor and recipient outcomes:

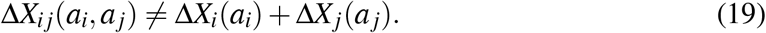

### Failure decomposition

A major advantage of the model is that the absence of recipient-benefiting action can be decomposed into mechanistically different failures. The decomposition follows the components defined above: inferred recipient need 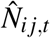, affordance recognition *Q*_*i j,t*_(*a*), valuation after cost, competition, relationship, solicitation, joint-payoff, and bias terms *U*_*i,t*_(*a*), and final action selection *P*_*i*_(*a* | *z*_*t*_). These components are derived from the independently specified task and social variables in Table 2. They are not inferred from whether recipient-benefiting action occurred. For this decomposition, continuous model quantities are summarised by binary diagnostic indicators. The indicator 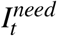is derived from the inferred-need estimate 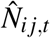and records whether recipient need was sufficiently inferred. The indicator 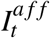is derived from the affordance-recognition term *Q*_*i j,t*_(*a*) and records whether an effective recipient-benefiting action was represented. The indicator 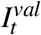 is derived from the utility term *U*_*i,t*_(*a*) and records whether the recipient-benefiting action retained positive value after cost, competition, relationship, solicitation, joint payoff, and bias terms were evaluated. The indicator 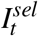 is derived from the softmax choice probability *P*_*i*_(*a* | *z*_*t*_) and records whether the recipient-benefiting action was selected. These indicators are introduced only for diagnostic decomposition. They are not additional latent mechanisms. The probability that all components align to produce recipient-benefiting action can then be written as:

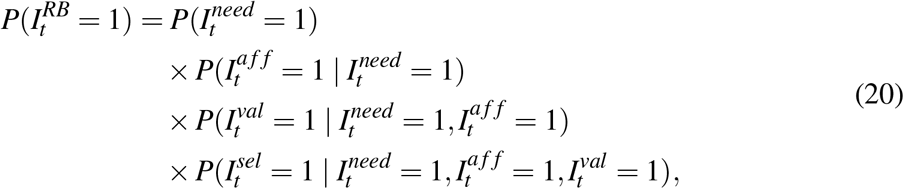

where 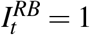 denotes the realised occurrence of recipient-benefiting action. The expression is a diagnostic chain: recipient need must be inferred, an effective action must be recognised, valuation must favour the action after costs and social terms are evaluated, and the action must then be selected. The realised helping label is assigned only after the selected action has changed, or failed to change, the recipient’s state:

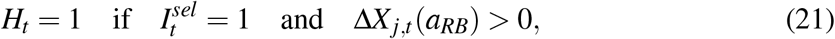

where *a*_*RB*_ denotes the action that would benefit the recipient in the relevant task. Thus, a trial can fail to produce helping because need was not inferred, because no effective affordance was recognised, because valuation suppressed the action, because stochastic action selection favoured another option, or because a help-like action was selected but did not actually benefit the recipient. The latter case is the false-positive condition. A null result is therefore underidentified unless the relevant part of the process can be constrained: need inference, affordance recognition, valuation, action selection, or realised recipient benefit. Table 3 lists the major failure classes.

**Table 3:**
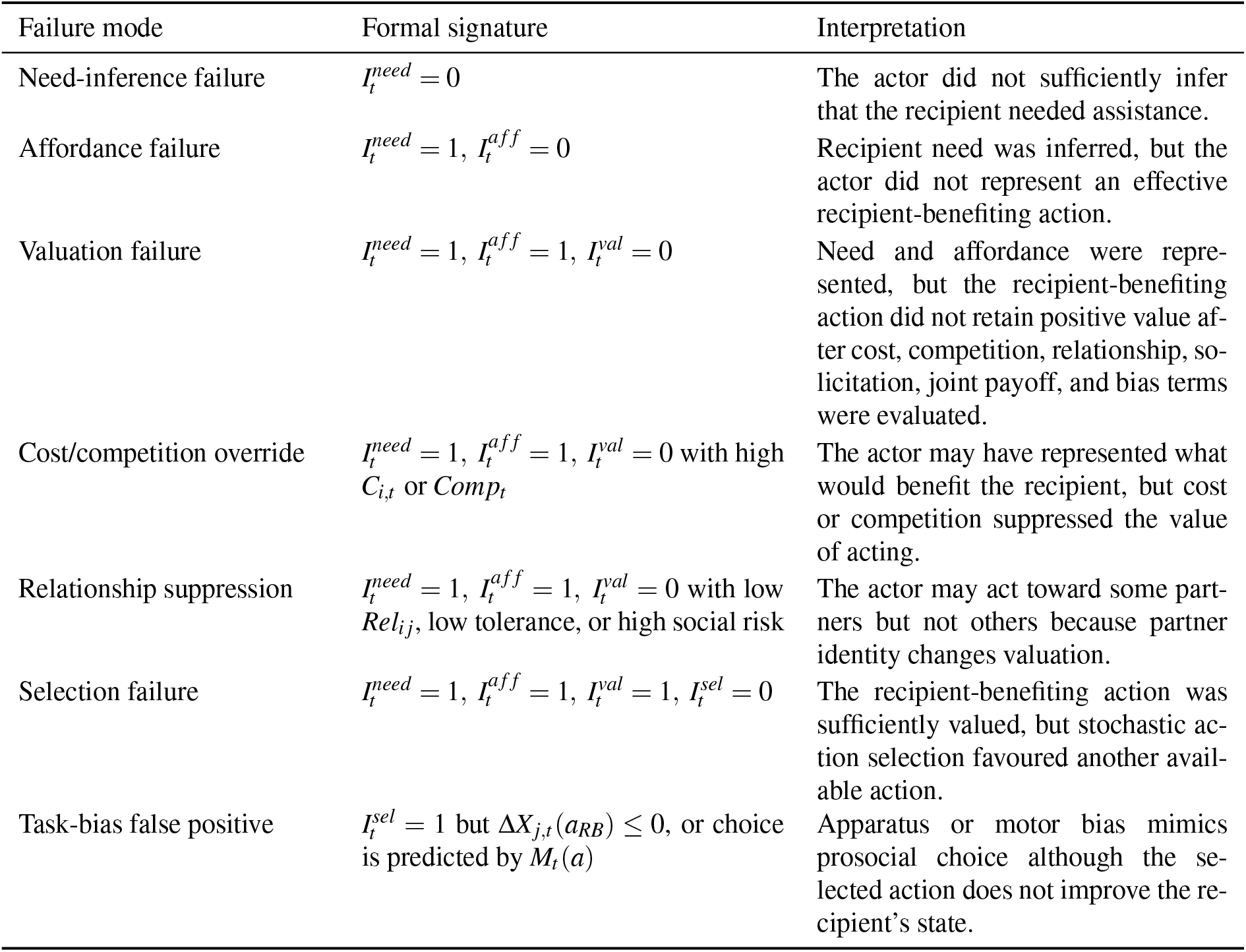
Failure modes that can all appear behaviourally as “no helping”.

### Task presets and diagnostic contrasts

Different experimental paradigms can be represented as different regions of the same input space. The simulation implemented ten task presets: instrumental reaching, visible tool transfer, hidden tool transfer, randomised food-token choice, fixed-position food-token choice, apparatus-bias/no-benefit trials, low-competition provisioning, high-cost helping, dominance-risk helping, and reliable-partner collaboration. The diagnostic logic of these presets is summarised in the Supplementary Material, Table S2. Table 4 translates the framework into diagnostic contrasts. The aim is not to assign a single psychological label to each paradigm, but to specify which alternative mechanisms must be separated before a helping outcome can support a stronger interpretation.

**Table 4:**
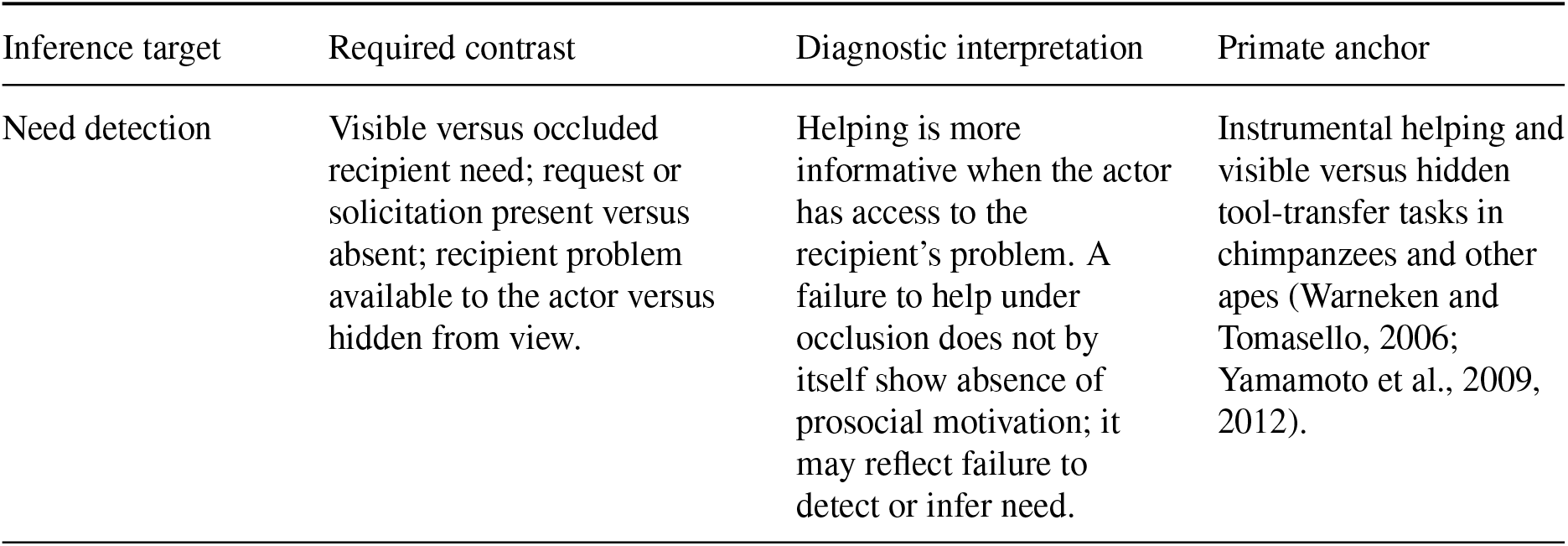
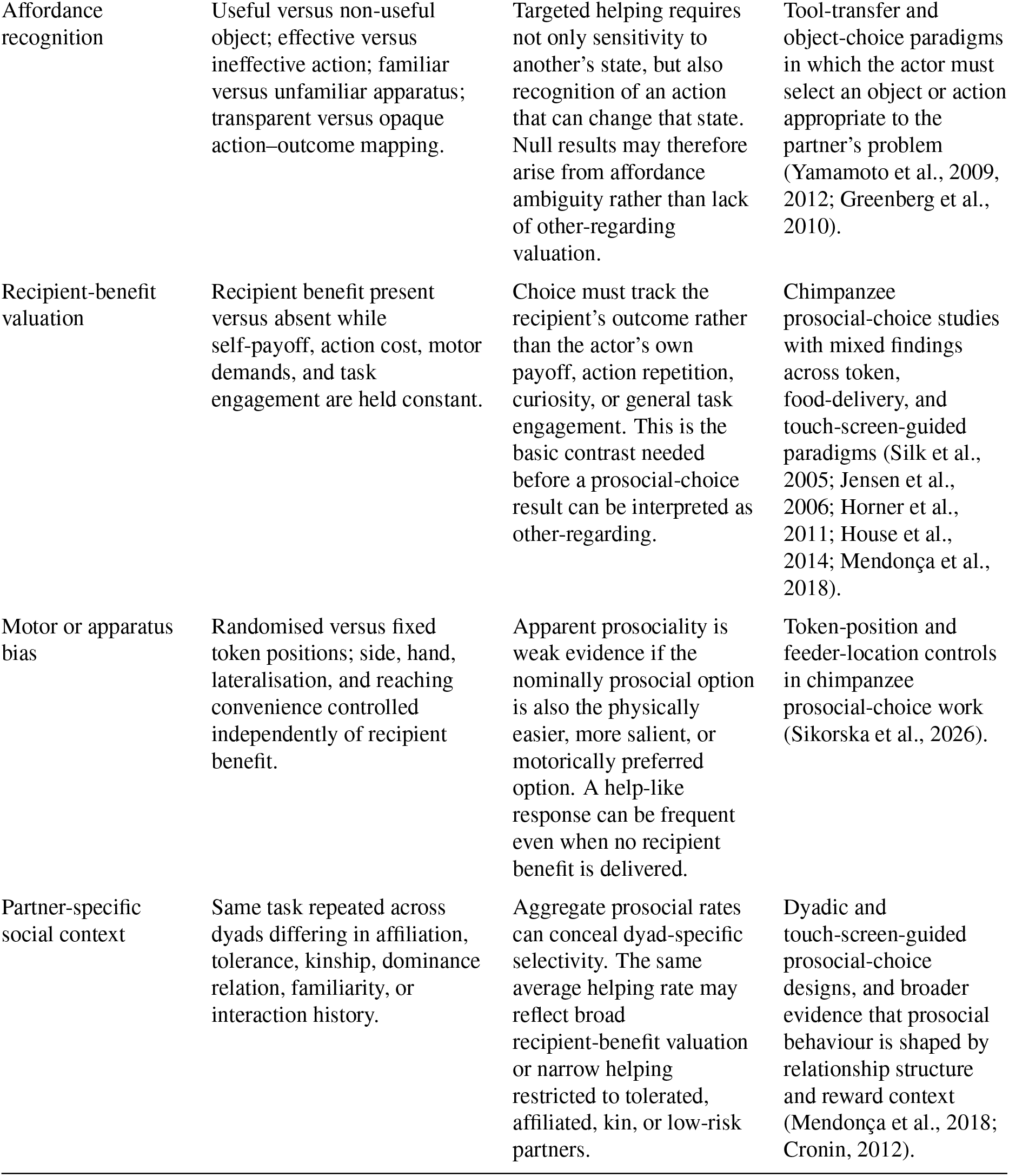
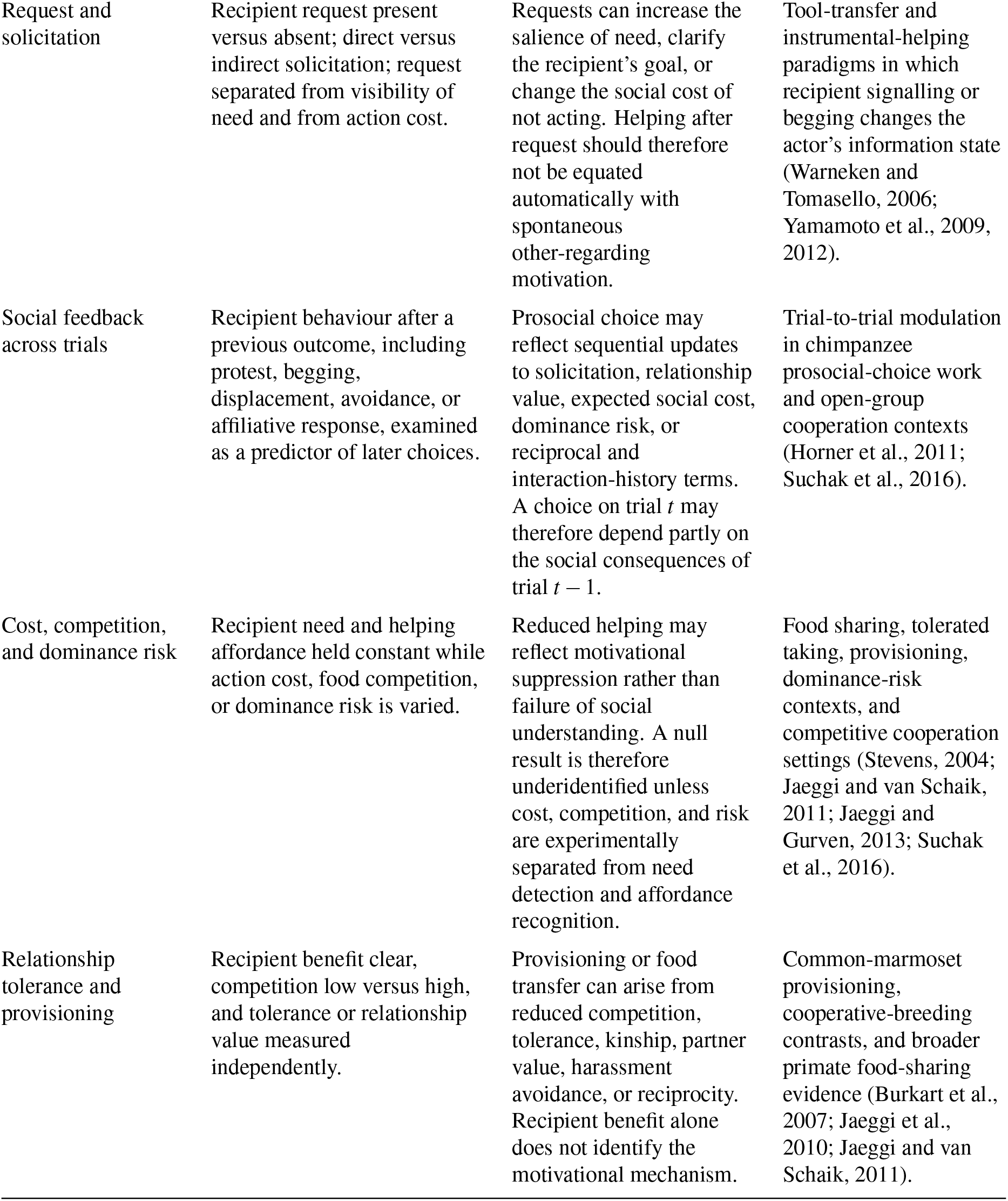
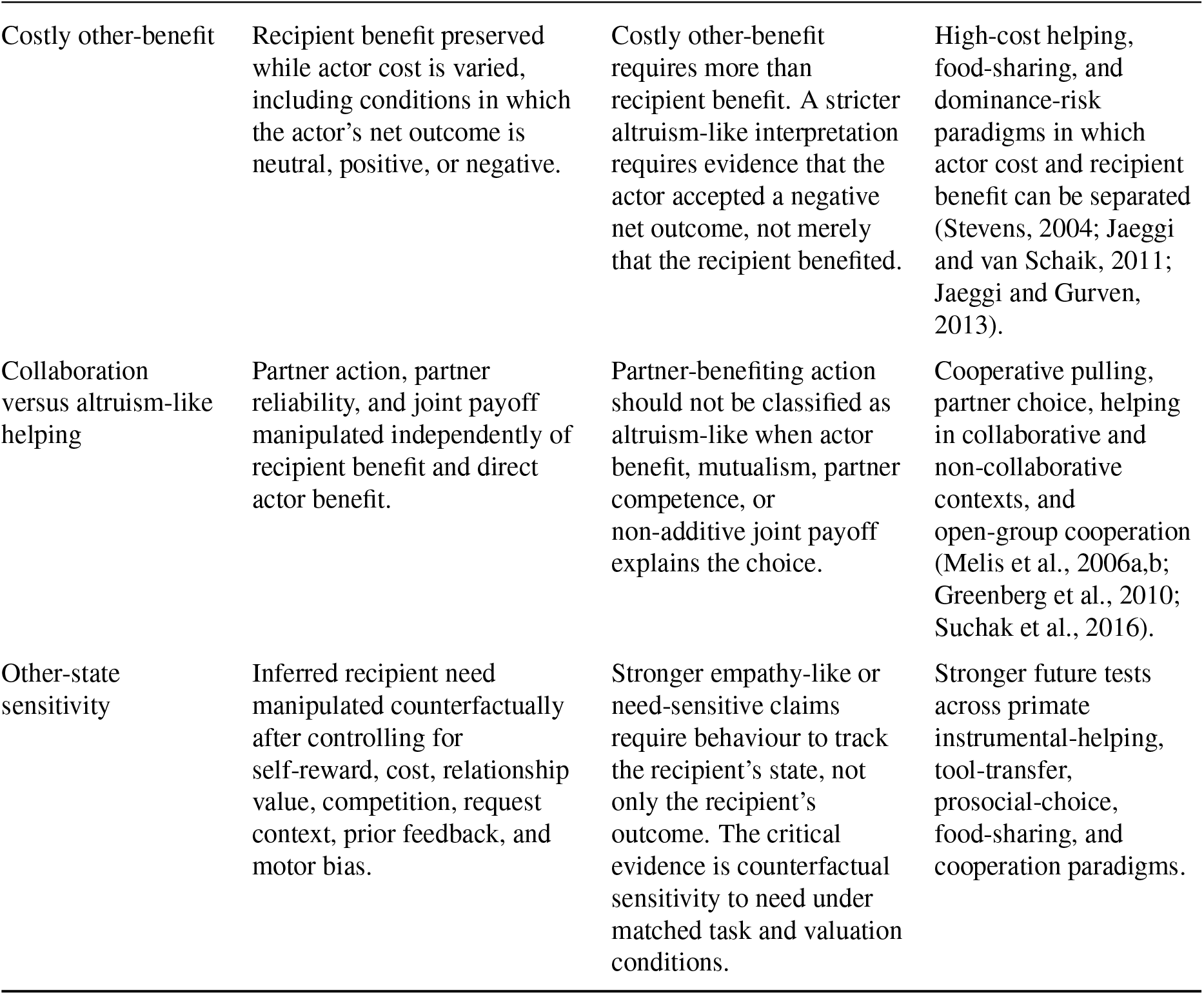
Diagnostic contrasts for interpreting recipient-benefiting outcomes in primate helping and prosocial-choice paradigms. The table specifies which alternative explanations must be separated before stronger claims about prosocial motivation, empathy-like responding, altruism-like action, or collaboration are justified.

| Inference target | Required contrast | Diagnostic interpretation | Primate anchor |
| --- | --- | --- | --- |
| Need detection | Visible versus occluded recipient need; request or solicitation present versus absent; recipient problem available to the actor versus hidden from view. | Helping is more informative when the actor has access to the recipient’s problem. A failure to help under occlusion does not by itself show absence of prosocial motivation; it may reflect failure to detect or infer need. | Instrumental helping and visible versus hidden tool-transfer tasks in chimpanzees and other apes (Warneken and Tomasello, 2006; Yamamoto et al., 2009, 2012). |
| Affordance recognition | Useful versus non-useful object; effective versus ineffective action; familiar versus unfamiliar apparatus; transparent versus opaque action–outcome mapping. | Targeted helping requires not only sensitivity to another’s state, but also recognition of an action that can change that state. Null results may therefore arise from affordance ambiguity rather than lack of other-regarding valuation. | Tool-transfer and object-choice paradigms in which the actor must select an object or action appropriate to the partner’s problem (Yamamoto et al., 2009, 2012; Greenberg et al., 2010). |
| Recipient-benefit valuation | Recipient benefit present versus absent while self-payoff, action cost, motor demands, and task engagement are held constant. | Choice must track the recipient’s outcome rather than the actor’s own payoff, action repetition, curiosity, or general task engagement. This is the basic contrast needed before a prosocial-choice result can be interpreted as other-regarding. | Chimpanzee prosocial-choice studies with mixed findings across token, food-delivery, and touch-screen-guided paradigms (Silk et al., 2005; Jensen et al., 2006; Horner et al., 2011; House et al., 2014; Mendonça et al., 2018). |
| Motor or apparatus bias | Randomised versus fixed token positions; side, hand, lateralisation, and reaching convenience controlled independently of recipient benefit. | Apparent prosociality is weak evidence if the nominally prosocial option is also the physically easier, more salient, or motorically preferred option. A help-like response can be frequent even when no recipient benefit is delivered. | Token-position and feeder-location controls in chimpanzee prosocial-choice work (Sikorska et al., 2026). |
| Partner-specific social context | Same task repeated across dyads differing in affiliation, tolerance, kinship, dominance relation, familiarity, or interaction history. | Aggregate prosocial rates can conceal dyad-specific selectivity. The same average helping rate may reflect broad recipient-benefit valuation or narrow helping restricted to tolerated, affiliated, kin, or low-risk partners. | Dyadic and touch-screen-guided prosocial-choice designs, and broader evidence that prosocial behaviour is shaped by relationship structure and reward context (Mendonça et al., 2018; Cronin, 2012). |
| Request and solicitation | Recipient request present versus absent; direct versus indirect solicitation; request separated from visibility of need and from action cost. | Requests can increase the salience of need, clarify the recipient's goal, or change the social cost of not acting. Helping after request should therefore not be equated automatically with spontaneous other-regarding motivation. | Tool-transfer and instrumental-helping paradigms in which recipient signalling or begging changes the actor's information state (Warneken and Tomasello, 2006; Yamamoto et al., 2009, 2012). |
| Social feedback across trials | Recipient behaviour after a previous outcome, including protest, begging, displacement, avoidance, or affiliative response, examined as a predictor of later choices. | Prosocial choice may reflect sequential updates to solicitation, relationship value, expected social cost, dominance risk, or reciprocal and interaction-history terms. A choice on trial $t$ may therefore depend partly on the social consequences of trial $t - 1$. | Trial-to-trial modulation in chimpanzee prosocial-choice work and open-group cooperation contexts (Horner et al., 2011; Suchak et al., 2016). |
| Cost, competition, and dominance risk | Recipient need and helping affordance held constant while action cost, food competition, or dominance risk is varied. | Reduced helping may reflect motivational suppression rather than failure of social understanding. A null result is therefore underidentified unless cost, competition, and risk are experimentally separated from need detection and affordance recognition. | Food sharing, tolerated taking, provisioning, dominance-risk contexts, and competitive cooperation settings (Stevens, 2004; Jaeggi and van Schaik, 2011; Jaeggi and Gurven, 2013; Suchak et al., 2016). |
| Relationship tolerance and provisioning | Recipient benefit clear, competition low versus high, and tolerance or relationship value measured independently. | Provisioning or food transfer can arise from reduced competition, tolerance, kinship, partner value, harassment avoidance, or reciprocity. Recipient benefit alone does not identify the motivational mechanism. | Common-marmoset provisioning, cooperative-breeding contrasts, and broader primate food-sharing evidence (Burkart et al., 2007; Jaeggi et al., 2010; Jaeggi and van Schaik, 2011). |
| Costly other-benefit | Recipient benefit preserved while actor cost is varied, including conditions in which the actor's net outcome is neutral, positive, or negative. | Costly other-benefit requires more than recipient benefit. A stricter altruism-like interpretation requires evidence that the actor accepted a negative net outcome, not merely that the recipient benefited. | High-cost helping, food-sharing, and dominance-risk paradigms in which actor cost and recipient benefit can be separated (Stevens, 2004; Jaeggi and van Schaik, 2011; Jaeggi and Gurven, 2013). |
| Collaboration versus altruism-like helping | Partner action, partner reliability, and joint payoff manipulated independently of recipient benefit and direct actor benefit. | Partner-benefiting action should not be classified as altruism-like when actor benefit, mutualism, partner competence, or non-additive joint payoff explains the choice. | Cooperative pulling, partner choice, helping in collaborative and non-collaborative contexts, and open-group cooperation (Melis et al., 2006a,b; Greenberg et al., 2010; Suchak et al., 2016). |
| Other-state sensitivity | Inferred recipient need manipulated counterfactually after controlling for self-reward, cost, relationship value, competition, request context, prior feedback, and motor bias. | Stronger empathy-like or need-sensitive claims require behaviour to track the recipient's state, not only the recipient's outcome. The critical evidence is counterfactual sensitivity to need under matched task and valuation conditions. | Stronger future tests across primate instrumental-helping, tool-transfer, prosocial-choice, food-sharing, and cooperation paradigms. |

### Simulation design: task presets and parameter presets

The accompanying MATLAB simulation implements a minimal two-action version of the model. On each trial, the actor chooses between a recipient-benefiting action and a nonrecipient-benefiting action. The simulation is organised as a crossing of task presets and parameter presets. Task presets define the externally specified problem structure of the simulated paradigm. Parameter presets define the actor-side valuation structure applied to that problem. A reported helping rate therefore always refers to one task preset evaluated under one parameter preset.

Task presets specify the task context before action selection occurs. They set variables such as visibility, solicitation, goal transparency, affordance clarity, familiarity, occlusion, action cost, competition, relationship value, reciprocal history, direct actor benefit, recipient benefit, joint benefit, and motor or apparatus bias. These variables correspond to the model inputs in Table 2, including *V*_*t*_, *S*_*t*_, *G*_*t*_, *A*_*t*_(*a*), *F*_*t*_, *O*_*t*_, *C*_*i,t*_(*a*), *Comp*_*t*_, *Rel*_*i j*_, *Recip*_*i j*_, 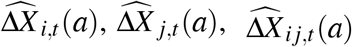, and *M*_*t*_(*a*). For example, a low-competition task preset means that the simulated task has a low value of *Comp*_*t*_. It does not imply that the simulated actor is weakly sensitive to competition. That sensitivity is defined by the parameter preset. The task presets used in the illustrative simulation are summarised in Table 5. These presets were not fitted to empirical data and should not be interpreted as species-specific paradigms. Instead, they define theoretically relevant assay structures in which visibility, solicitation, goal transparency, affordance clarity, occlusion, cost, competition, relationship value, reciprocity, direct actor benefit, recipient benefit, joint benefit, and motor or apparatus bias can be independently varied. The full numerical task-preset matrix used in the MATLAB implementation is provided in the Supplementary Material, Table S4.

**Table 5:** Task presets used in the illustrative simulation. Each preset specifies the external task structure before action selection occurs. The values listed here describe the intended model contrast; the full numerical task-preset matrix is provided in the Supplementary Material, Table S4.

| Task preset | Assay structure | Main diagnostic contrast |
| --- | --- | --- |
| Instrumental reaching | Highly visible, solicited, goal-transparent, low-cost helping opportunity with clear action affordance and low competition. | Tests whether a transparent recipient-only helping opportunity is selected when need and affordance information are available. |
| Visible tool transfer | Visible transfer of an object or tool that benefits the recipient, with high goal transparency, high affordance clarity, low occlusion, low cost, and low competition. | Tests recipient-benefiting action under a clear instrumental structure without direct actor benefit. |
| Hidden tool transfer | Tool-transfer structure with reduced visibility, lower goal transparency, weaker affordance clarity, and high occlusion. | Tests whether recipient-benefiting action is suppressed when the same possible benefit is less perceptually and causally available to the actor. |
| Food-token randomised | Abstract token-choice structure with weak solicitation, moderate goal transparency, low cost, and relatively high competition. | Tests whether recipient benefit is selected in a less transparent, more abstract, and more competitive task context. |
| Food-token fixed position | Token-choice structure similar to the randomised food-token preset, but with an added fixed-position or motor bias against the recipient-benefiting option. | Tests whether apparatus or side biases can reduce apparent helping independently of recipient benefit. |
| Apparatus bias, no benefit | Low-cost task with no recipient benefit, no relationship or reciprocity value, and a strong motor or apparatus bias toward the nominal help response. | Tests false-positive helping: a help-like action can be selected even though the recipient's state is not improved. |
| Low-competition provisioning | Moderately transparent provisioning opportunity with low cost, low competition, moderate relationship value, and recipient-only benefit. | Tests recipient-benefiting action when competitive suppression is weak and the task is less abstract than token choice. |
| High-cost help | Transparent, solicited helping opportunity with high recipient benefit but high actor cost. | Tests whether recipient-benefiting action is suppressed by cost even when need and affordance information are clear. |
| Dominance-risk help | Transparent helping opportunity with moderate-to-high cost and high competition or dominance risk. | Tests whether competitive or dominance-related suppression prevents helping despite visible recipient benefit. |
| Reliable-partner collaboration | Collaborative task in which the selected action benefits the actor, benefits the recipient, and produces an additional joint payoff. | Tests whether action can be explained by direct actor benefit and joint payoff, rather than by recipient-only benefit. |

Parameter presets specify how strongly the simulated actor weights task variables during valuation. They set weights such as *w*_*self*_, *w*_*other*_, *w*_*cost*_, *w*_*comp*_, *w*_*rel*_, *w*_*recip*_, *w*_*sol*_, and *w*_*joint*_ in Equation 9. For example, a high-competition-sensitivity parameter preset means that *w*_*comp*_ is high, so competition strongly suppresses the value of acting. A low-competition-sensitivity parameter preset means that *w*_*comp*_ is low, so the same competitive task context has a weaker suppressive effect. The parameter *w*_*other*_ sets the baseline weight assigned to the recipient’s expected outcome 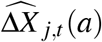 once need inference and affordance recognition have opened the social gate. Parameter presets with larger values of *w*_*other*_ are therefore described as having stronger other-outcome weighting. This label should not be read as altruism, empathy, or prosocial motivation. It only specifies how strongly recipient benefit contributes to utility after the relevant need and affordance have been represented.

This distinction between task presets and parameter presets is central to the interpretation of the simulation. Task presets define the external assay. Parameter presets define the actor configuration tested within that assay. The same task preset can therefore produce different levels of recipient-benefiting action under different parameter presets. Conversely, the same parameter preset can produce different outcomes across task presets when visibility, cost, competition, actor benefit, recipient benefit, joint payoff, or apparatus bias are changed. The simulation is therefore not designed to assign a fixed psychological interpretation to any particular task. It is designed to show how task structure and valuation parameters jointly determine realised helping. The simulation follows the formal distinction between direct actor benefit, recipient benefit, and joint benefit introduced above. In non-collaborative recipient-benefiting task presets, the recipient-benefiting action can improve the recipient’s state while providing no direct actor benefit. In the collaboration task preset, coordinated action can improve both the actor’s and the recipient’s outcomes, and may also produce a joint benefit. This distinction is important because self-interest, recipient-outcome sensitivity, and joint-payoff sensitivity are not treated as interchangeable. A self-interest control can explain collaborative action only when the selected action also improves the actor’s own outcome. It cannot explain recipient-only benefit by reclassifying recipient benefit as actor benefit. The simulation also records the stage at which recipient-benefiting action fails. A non-recipient-benefiting outcome can therefore be classified as need-inference failure, affordance failure, valuation failure, selection failure, or realised-outcome failure. Realised-outcome failure corresponds to a false-positive case: a help-like action is selected, for example because of motor or apparatus bias, but the recipient’s state is not improved. The simulation is therefore useful not only for generating recipient-benefiting action, but also for showing why both positive and null findings can be ambiguous.

All simulations used 500 trials per task preset and parameter preset. The random seed was fixed in the MATLAB reproduction script for the illustrative run reported here. Choice was stochastic and followed the softmax rule in Equation 10. Unless otherwise stated, the utility of the non-recipient-benefiting action was set to zero. Action initiation was governed by a task-independent threshold, with a lower threshold when direct actor benefit was present. The recipient-benefiting action threshold was set to 0.25, the actor-benefit action threshold to 0.05, and the action-benefit tolerance to 10^−9^. This preserves stochastic low-cost action while preventing recipient-only benefit from being counted as direct actor benefit.

The parameter presets used in the simulation are not intended as direct species models. They represent theoretically relevant regions of model space. For example, a moderate-other/highcompetition-sensitivity preset captures a system in which recipient outcomes can affect choice but are easily suppressed by resource competition. A high-other/low-competition-sensitivity preset captures a system in which recipient outcomes receive stronger weight and competitive suppression is weaker. A low-other-weight preset captures weak sensitivity to recipient benefit, whereas a pure self-interest control removes other-outcome, relationship, reciprocity, solicitation, and joint-payoff terms. Species-level hypotheses should therefore be expressed as predictions about parameter distributions, not as fixed labels attached to taxa. The parameter presets are summarised in Table 6. Full supplementary documentation is provided in the Supplementary Material: task-preset fields in Table S3, the full task-preset matrix in Table S4, parameter-preset fields in Table S5, the full parameter-preset matrix in Table S6, and output-summary columns in Table S7.

**Table 6:** Parameter presets used in the illustrative simulation. These presets define interpretable regions of parameter space and are not species simulations.

| Parameter preset | Main parameter emphasis | Interpretation |
| --- | --- | --- |
| Moderate other-weight / high competition sensitivity | $w_{other} = 0.28$, $w_{comp} = 1.05$,<br>$w_{cost} = 1.35$ | Recipient outcome can influence action valuation, but helping is strongly suppressed by competition and cost. |
| High other-weight / moderate competition sensitivity | $w_{other} = 0.95$, $w_{comp} = 0.55$,<br>$w_{cost} = 1.00$ | Recipient outcome has a strong effect on valuation, while competition and cost still constrain action selection. |
| High other-weight / low competition sensitivity | $w_{other} = 0.65$, $w_{comp} = 0.45$,<br>$w_{cost} = 1.15$ | Recipient outcome is weighted substantially and competitive suppression is comparatively weak. |
| Low other-weight | $w_{other} = 0.05$, $w_{comp} = 1.10$,<br>$w_{cost} = 1.60$ | Recipient outcome has little direct effect on valuation; low-cost or biased responses may still occur through residual non-recipient terms. |
| Pure self-interest | $w_{other} = w_{joint} = w_{rel} = w_{recip} =$<br>$w_{sol} = 0$ | Recipient outcome, joint payoff, relationship value, reciprocity, and solicitation are removed from valuation. |

## Model-ablation scoring

Recipient-benefiting action should not be interpreted by default as evidence that the actor valued the recipient’s outcome. Other-regarding valuation is one possible explanation, but it must be distinguished from low cost, direct actor benefit, joint payoff, relationship effects, solicitation, competition, and motor or apparatus bias. This issue was examined with model-ablation scoring. The ablation analysis is not presented as formal empirical model selection, because the data are generated by the full architecture and are not fitted to independent animal behaviour. Instead, the analysis asks how much predictive adequacy is lost when key components of the architecture are removed or simplified. Model-ablation scoring was therefore used as an internal diagnostic check. The question was whether the choices generated by the full social-gating model could also be explained by simpler utility functions. Each ablation keeps only a restricted subset of the terms in Equation 9. If a reduced model scores well, the corresponding behaviour does not require the full social-gating interpretation. If the full model scores better, this indicates that need inference, affordance recognition, recipient benefit, relationship modulation, solicitation, or joint payoff contributed information that the simpler model could not capture.

Bias-only ablation:

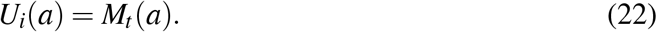

This model assumes that choice is explained only by motor or apparatus bias. It is a control for cases in which the nominally prosocial option is selected because it is easier, more salient, or spatially favoured, not because it benefits the recipient.

Self-interest ablation:

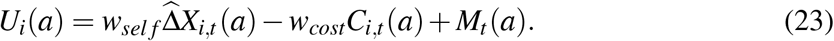

This model allows direct actor benefit and action cost, but contains no recipient-benefit term. It asks whether the generated choices can be explained as ordinary self-interested action plus motor or apparatus bias.

Competition ablation:

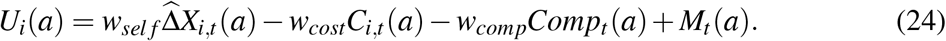

This model adds the suppressive effect of competition. It asks whether apparent non-helping can be explained by food competition or dominance-related inhibition without invoking absence of social understanding or other-regarding valuation.

Relationship ablation:

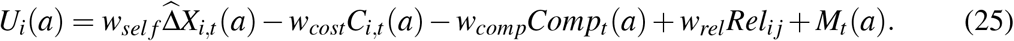

This model adds partner relationship value, but still omits the full social gate. It asks whether partner-specific valuation alone can account for the generated choices, without requiring explicit need detection, affordance recognition, or recipient-benefit sensitivity.

Full social-gating model: The full model is Equation 9. It includes direct actor benefit, cost, competition, motor or apparatus bias, socially gated recipient benefit, relationship modulation, reciprocity, solicitation, and joint payoff.

For simulated actions *a*_*1*_, …, *a*_*T*_, each ablation was scored by the log likelihood of the generated choices:

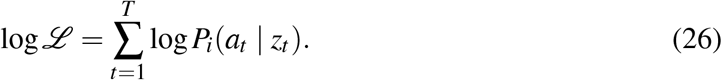

The log likelihood measures how well each reduced model predicts the actions generated in the simulation. A higher value means that the model assigned higher probability to the choices that actually occurred. The ablation analysis is therefore not an empirical model comparison across real animals; it is an internal check of whether the simulated behaviour depends on the components that the full architecture claims to separate. Scores are therefore used only as a component-ablation check, indicating which parts of the architecture are needed to reproduce the simulated choice patterns.

## Results

The results should be interpreted as a generative proof of principle rather than as fitted empirical estimates. The aim was to test whether one socially gated action-selection architecture can generate successes, failures, false negatives, and false positives under independently specified task conditions. The resulting overview of realised helping and counterfactual need sensitivity is shown in Figure 1.

**Figure 1:**
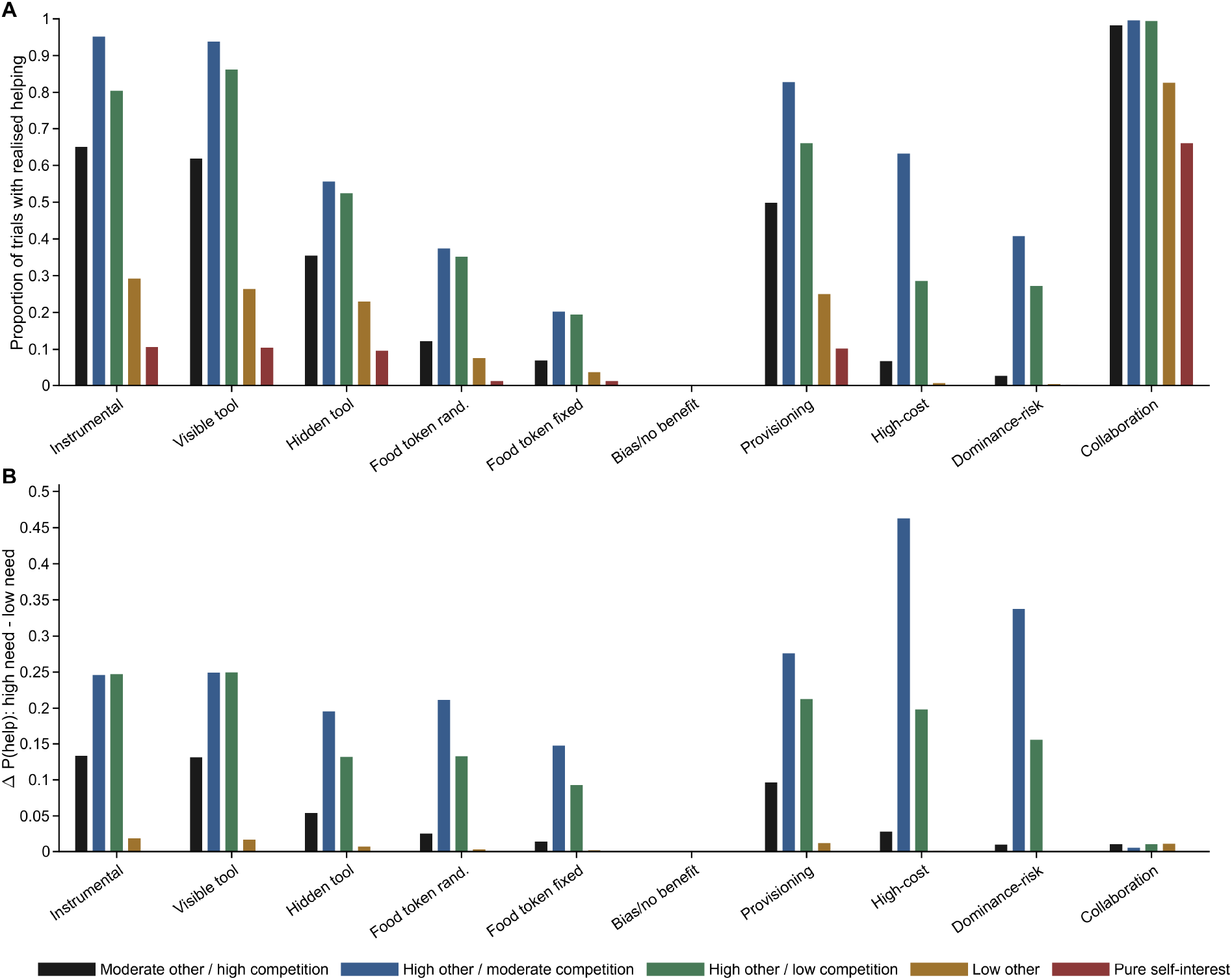
Realised helping and counterfactual need sensitivity across task presets and parameter presets. (A) Proportion of trials in which the selected action improved the recipient’s state. (B) Counterfactual need sensitivity, defined as the change in helping probability when inferred recipient need was increased while other task variables were held constant. Transparent instrumental and visible tool-transfer presets produced high realised helping when social valuation was present, whereas food-token, high-cost, and dominance-risk presets were more strongly constrained by task structure, competition, or cost. The collaboration preset remains high even in the pure self-interest control because coordinated action provides direct actor benefit and joint payoff.

### Task-dependent prosocial outcomes

Realised helping depended jointly on task structure and parameter preset. The same task could generate different helping rates under different valuation settings, and the same valuation setting produced different outcomes across task contexts. The strongest helping rates occurred in transparent, low-cost tasks when other-outcome weighting was moderate or high. Instrumental reaching produced helping rates of 0.65 under the moderate-other/high-competition preset, 0.95 under the high-other/moderate-competition preset, and 0.80 under the high-other/low-competition preset. Visible tool transfer showed the same qualitative pattern, with helping rates of 0.62, 0.94, and 0.86, respectively. The pure self-interest control helped less often in these non-collaborative tasks (0.11 for instrumental reaching and 0.10 for visible tool transfer), because recipient benefit no longer entered the actor’s self-benefit term. Residual low-cost helping in the pure self-interest control is not interpreted as social valuation. It follows from stochastic softmax action selection when the action is transparent and cheap. The diagnostic issue is therefore not whether recipient-only helping is exactly zero in every low-cost context, but whether it scales with inferred recipient need, survives cost and risk, and dissociates from apparatus bias. Occluding the recipient’s problem reduced helping. Hidden tool transfer produced lower helping than visible tool transfer across parameter presets, consistent with a failure at the need-detection or affordance-recognition stage. Food-token tasks were weaker and more parameter-sensitive. Helping rates in the randomised food-token task ranged from 0.01 in the pure self-interest control to 0.37 under the high-other/moderate-competition preset. Fixed-position food-token trials were lower still, with a maximum helping rate of 0.20, consistent with the idea that abstract food delivery is a poor assay unless side bias, motor preference, and action-outcome understanding are controlled. These task- and parameter-dependent patterns are summarised in Figure 1.

### Cost, risk, and collaboration

High-cost and dominance-risk tasks created a different type of failure. In these tasks, need and affordance were transparent, but the helping action was costly. Helping in the high-cost task was absent in the pure self-interest control and very low under the low-other-weight preset (0.006), but rose to 0.63 in the high-other/moderate-competition preset. Dominance-risk helping showed the same qualitative pattern: helping was 0.002 in the pure self-interest control and 0.41 in the high-other/moderate-competition preset. These cases therefore represent motivational suppression rather than failure to detect need. Reliable-partner collaboration produced high action rates under parameter presets with social valuation and remained substantial in the pure self-interest control (0.66). This is not a failure of the control condition. It follows from the task definition: collaboration carries direct actor benefit and joint payoff. Collaboration should therefore not be interpreted as altruism-like merely because the partner benefits. It is a distinct action context in which actor and recipient outcomes are coupled.

### False-positive prosociality

The apparatus-bias/no-benefit condition provides the clearest demonstration of why prosocial-choice tasks can be misleading. In this condition, the help-like option was favoured by motor or apparatus bias, but it produced no recipient benefit. The model nevertheless selected the help-like option at rates between 0.994 and 1.000 across parameter presets. Actual helping, prosocial choice, costly other-benefit, and strict altruism-like outcomes were all zero. Thus, choice of the nominally prosocial option and actual improvement of the recipient’s state were completely dissociated. This simulated dissociation captures the same methodological problem highlighted by recent chimpanzee token-choice work: when token placement, reaching convenience, and motor lateralisation are insufficiently controlled, choices may track the physical action structure rather than the social consequence of the selected option (Sikorska et al., 2026). The diagnostic requirement is therefore not simply to record which token or option was chosen, but to show that choice tracks recipient benefit after side, hand, location, and physical convenience have been controlled. A compact supplementary description of this false-positive apparatus-bias check is provided in the Supplementary Material.

### Failure decomposition

Failure decomposition showed that non-helping is not a single phenomenon. In instrumental reaching and visible tool transfer, failures were mainly valuation failures, because need and affordance were generally available. In hidden tool transfer, need-detection and affordance failures increased. In food-token tasks, non-helping reflected a mixture of need-detection failure, affordance ambiguity, weak valuation, and cost or competition. In high-cost and dominance-risk tasks, cost/competition override dominated. The apparatus-bias/no-benefit condition was a pure false positive: the help-like action was frequently selected, but recipient benefit was absent. The averaged decomposition is shown in Figure 2; the same decomposition separated by parameter preset is provided in the Supplementary Material, Fig. S1.

**Figure 2:**
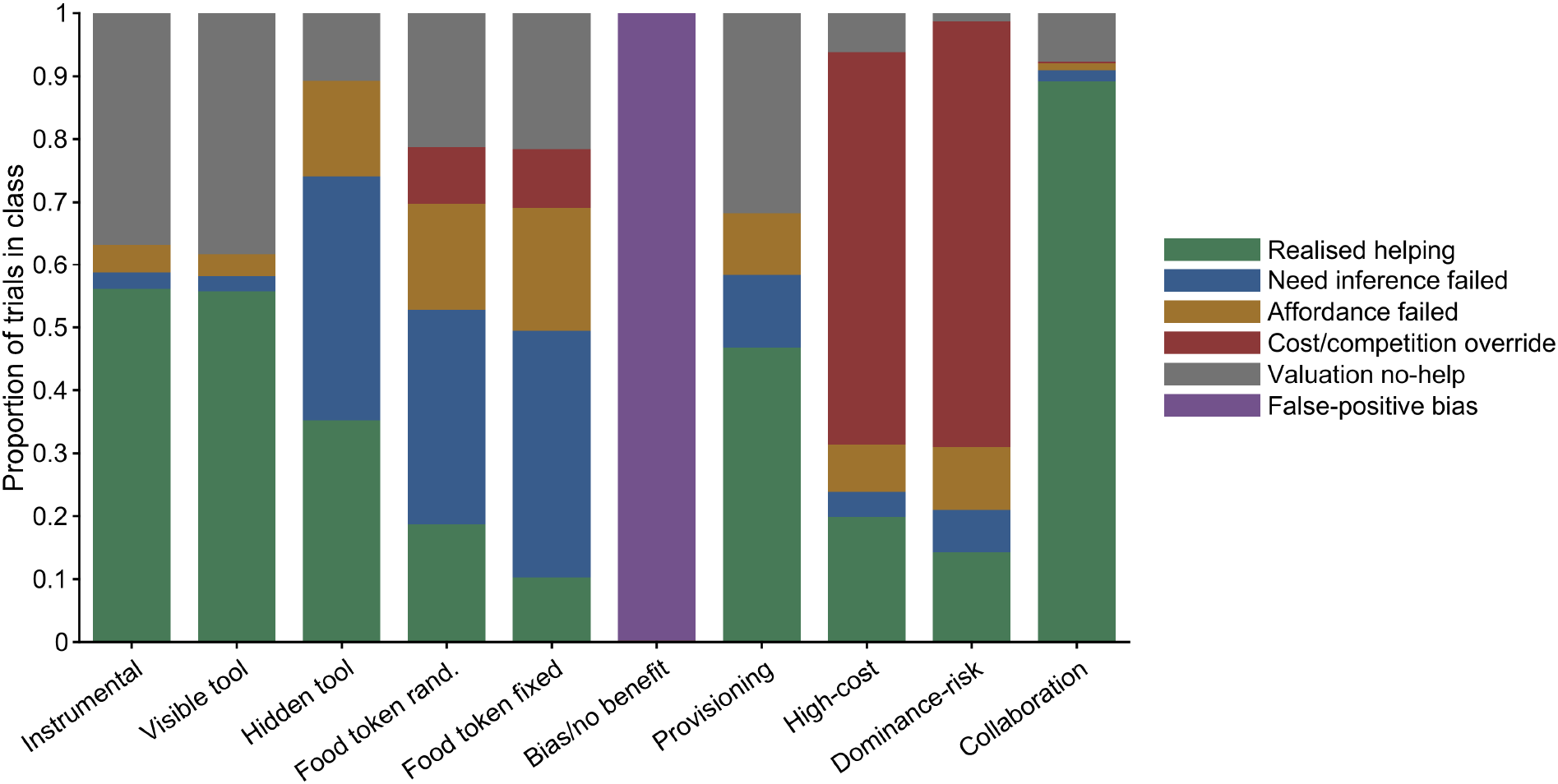
Failure decomposition by task preset, averaged across parameter presets. Bars show the proportion of trials assigned to each success or failure class. The same absence of realised helping can arise from different sources, including need-inference failure, affordance failure, cost or competition override, valuation-based non-helping, or false-positive apparatus bias. This decomposition illustrates why null results in primate helping and prosocial-choice tasks are underidentified unless the relevant task, inference, and valuation components are separated. The collaboration preset is included as a joint-payoff case rather than as recipient-only helping, illustrating why partner-benefiting action should not automatically be classified as altruism-like.

### Counterfactual need sensitivity

Counterfactual need sensitivity provides a direct diagnostic of other-state-sensitive action. In the apparatus-bias/no-benefit condition, increasing inferred recipient need had no effect on helping probability. The pure self-interest control was also flat across tasks, as expected. In contrast, presets with high other-outcome weighting showed positive need sensitivity. Under the high-other/moderate-competition preset, the need-only counterfactual effect was 0.46 in the high-cost task and 0.34 in the dominance-risk task. Thus, the model separates bias-driven action from action that is genuinely sensitive to inferred recipient need.

### Ablation-score diagnostics

The component-ablation check asked whether the simulated choices required the full social-gating architecture, or whether they could be reproduced by simpler explanations such as motor bias, self-interest, competition, or relationship value alone. Because the choices were generated by the simulation rather than fitted to independent animal behaviour, this analysis should not be interpreted as empirical model validation. It instead asks whether the simulated output patterns depend on the components that the model is intended to separate.

The results showed the expected separation. In presets with nonzero other-outcome weighting, ablations that removed social-gating terms had poorer predictive scores, indicating that simpler utility functions could not fully reproduce the generated choice patterns. In the pure self-interest control, by contrast, simpler non-social ablations performed similarly or better than the full social-gating model. This is the appropriate outcome: when choices are generated without social valuation terms, the scoring procedure should not force an other-regarding explanation. Thus, the ablation scores served as an internal diagnostic check, distinguishing cases in which recipient-benefiting action depended on socially gated valuation from cases in which it could be explained by simpler controls. A compact supplementary description of the component-ablation check is provided in the Supplementary Material.

### Summary of simulation results

Taken together, the simulation produced four diagnostic signatures. First, transparent low-cost helping was high when need and affordance were available, but this did not require a dedicated helping mechanism distinct from ordinary action selection. Second, food-token and hidden-affordance tasks were weaker and more parameter-sensitive, showing how null or mixed findings can arise from uncertainty, competition, or abstract action–outcome mapping. Third, apparatus bias generated near-ceiling help-like choice without recipient benefit, demonstrating that apparent prosociality can be false-positive. Fourth, high-cost, dominance-risk, and collaboration presets dissociated recipient benefit from altruism-like interpretation: cost and risk suppressed helping even when need was represented, whereas collaboration remained possible under self-interest when actor benefit and joint payoff were present. These signatures motivate the literature-based interpretation developed in the Discussion.

## Discussion

The simulation results organise several otherwise difficult patterns in the primate helping and prosocial-choice literature. Transparent low-cost tasks produced high helping when recipient need and the effective action were available. This fits instrumental-helping and visible tool-transfer paradigms, where the recipient’s problem and the relevant affordance are comparatively clear (Warneken and Tomasello, 2006; Yamamoto et al., 2009, 2012). By contrast, hidden tool-transfer and food-token tasks were weaker and more parameter-sensitive, because they load simultaneously on need detection, action–outcome understanding, competition, partner context, and motor or apparatus bias. The main contribution of the model is therefore not to declare which studies show true prosocial motivation; it is to specify which latent components must be separated before such an interpretation is warranted.

### Organising the empirical literature

The simulation results should be read as a map of why apparently conflicting findings in primate helping research can all be valid within their own task contexts. Transparent low-cost tasks produced high helping rates when other-outcome weighting was moderate or high. This pattern fits the instrumental-helping and visible tool-transfer literatures, where recipient need, goal structure, and the effective action are relatively transparent (Warneken and Tomasello, 2006; Yamamoto et al., 2009, 2012). In model terms, such tasks reduce uncertainty at the need-detection and affordance-recognition stages. Positive performance in these contexts therefore demonstrates that the actor can act on another’s problem when the relevant information is available, but it does not by itself estimate a context-independent altruistic disposition. The weaker performance in hidden tool-transfer and food-token presets captures the complementary problem. In these tasks, the recipient’s need, the action–outcome mapping, or the social consequence of the action is more abstract. The model therefore predicts that helping should become more variable and more sensitive to solicitation, familiarity, competition, and task design. This provides a formal interpretation of why some prosocial-choice studies reported little or no other-regarding choice (Silk et al., 2005; Jensen et al., 2006), whereas other designs elicited stronger prosocial responding or revealed sensitivity to task structure and social context (House et al., 2014; Mendonça et al., 2018). The point is not that one set of findings is correct and the other is artefactual. The point is that food-token and apparatus-based paradigms load simultaneously on other-outcome valuation, action–outcome comprehension, food competition, partner context, and motor or side bias. The low-competition provisioning preset provides a separate interpretation of cooperative-breeding and food-sharing contrasts. Food transfer in primates can be shaped by tolerance, kinship, harassment, reciprocity, partner value, and ecological context rather than by one homogeneous prosocial motivation (Stevens, 2004; Jaeggi and van Schaik, 2011; Jaeggi and Gurven, 2013). Higher provisioning in such a context need not imply a qualitatively different moral faculty. In the model, it can emerge when recipient benefit is weighted more strongly, competition is weaker, or social tolerance reduces the effective cost of acting. This is consistent with treating marmoset provisioning and related cooperative-breeding findings as socio-ecologically tuned parameter configurations rather than as direct evidence for a unitary prosocial trait (Burkart et al., 2007; Jaeggi et al., 2010). The high-cost and dominance-risk presets show why null results are also underidentified. In those simulations, recipient need and helping affordance were available, but action was suppressed by cost, risk, or competition. This interpretation is consistent with food-sharing and cooperation work showing that recipient benefit can be constrained by harassment, tolerance, dominance, competition, partner value, and social enforcement (Stevens, 2004; Jaeggi and van Schaik, 2011; Jaeggi and Gurven, 2013; Suchak et al., 2016). A failure to help in such contexts should therefore not be interpreted as failure of social understanding unless need detection, affordance recognition, action cost, food competition, and social risk have been separated. Conversely, the reliable-partner collaboration preset shows why partner-benefiting action should not automatically be classified as altruism-like. Chimpanzee cooperation studies show that success can depend on tolerance constraints and on selecting competent partners (Melis et al., 2006a,b). In model terms, these effects belong to competition, relationship value, partner reliability, and joint-payoff terms rather than to recipient benefit alone. Open-group cooperation studies further show that cooperative success can be shaped by competition, displacement, freeloading, and social enforcement, which correspond in the present model to competition, dominance-risk, partner-history, and social-cost terms (Suchak et al., 2016). When actor benefit and joint payoff are present, the partner’s benefit is part of a coordinated payoff structure rather than evidence for costly other-regarding motivation.

### False-positive prosociality and task structure

The false-positive result is central for interpreting prosocial-choice tasks. In the simulation, the apparatus-bias/no-benefit preset produced near-ceiling help-like choice while actual recipient benefit remained absent. This is not a claim that published prosocial-choice effects are artefactual. Rather, it formalises a general inferential problem: a response can look prosocial at the level of task coding while being generated by motor convenience, side preference, token location, or apparatus structure. Sikorska et al. (Sikorska et al., 2026) provide a clear empirical example of this vulnerability in chimpanzees. When token placement reduced lateralisation effects, prosocial choices increased; when token positions were fixed, choices were systematically shaped by physical reaching convenience. In the present model, this ambiguity is represented by the motor/apparatus-bias term *M*_*t*_(*a*) and by the apparatus-bias/no-benefit task preset. Apparent prosociality therefore requires controls showing that choices track recipient outcomes rather than the physical affordances of the response.

### Social determinants of prosocial choice

The complementary problem is that genuine social modulation can also be missed if prosociality is treated as a single aggregate rate. Mendonça et al. (Mendonça et al., 2018) are important in this respect because their touch-screen-guided prosocial-choice design connects prosocial responding to the social context of the actor–recipient dyad. In the present framework, such effects are not treated as noise around a species-level prosocial tendency. They are represented explicitly as relationship value, reciprocal or interaction history, dominance risk, tolerance, competition, solicitation, and partner-specific valuation terms, consistent with broader evidence that prosocial behaviour is shaped by relationship structure, communication, and reward context (Cronin, 2012). The altruistic condition in Mendonça et al. (Mendonça et al., 2018) is particularly informative for the present framework because the active chimpanzee could deliver food to the recipient without obtaining food herself. Such trials separate recipient benefit from direct actor benefit more cleanly than mutualistic food-delivery conditions. The reported cases in which young female offspring acted in ways that benefited their mothers are therefore not merely additional prosocial choices; they point to a specific region of the model space in which recipient benefit, relationship value, tolerance, kinship, and low social risk may jointly overcome the absence of direct actor payoff. This is exactly the kind of pattern that should be analysed as a dyad- and context-specific parameter configuration rather than averaged into a species-level prosociality score. This matters because the same average prosocial rate can be produced by different mechanisms. One individual may show moderate helping because recipient benefit is weighted across many partners. Another may show the same average rate because helping is restricted to particular partners, such as tolerated, affiliated, kin, familiar, or low-risk recipients. Conversely, a low average prosocial rate may conceal selective helping toward a narrow subset of partners. The model therefore treats partner identity and dyadic context as explanatory variables rather than nuisance variation. Social determinants define the conditions under which detected recipient need and recognised helping affordance are allowed to influence action.

A further implication is that social determinants need not be static properties of the dyad. They can also operate dynamically across trials. Prosocial-choice work in chimpanzees suggests that recipient behaviour before or around a choice can modulate later prosocial responding (Horner et al., 2011). In the present framework, such effects are not treated as a separate psychological faculty. They can be represented as trial-to-trial updates to solicitation, relationship value, expected social cost, dominance risk, or reciprocal and interaction-history terms. In particular, the recipient’s reaction to a previous non-prosocial outcome can be interpreted as social feedback that changes the actor’s later valuation of the partner, the expected cost of not helping, or the salience of the recipient’s need. Different forms of protest need not have equivalent effects: direct protest may increase conflict, avoidance, or dominance-related cost, whereas indirect protest may function more like need signalling or solicitation. A choice on trial *t* may therefore reflect not only the current apparatus, but also the social consequences of trial *t* − 1. This trial-to-trial interpretation is useful because it prevents a misleading dichotomy between stable prosocial preference and immediate task bias. Prosocial action may depend on a temporally extended social process in which previous outcomes, recipient responses, protest, tolerance, and partner history alter the valuation structure of the next trial. Formally, such effects would be represented by allowing terms such as *Rel*_*i j*_, *Recip*_*i j*_, *S*_*t*_, and dominance or conflict-related costs to update across trials. The current simulation treats these terms as preset task and parameter values, but the same architecture could be extended to model sequential social feedback explicitly.

This interpretation also clarifies why social determinants and task determinants must be modelled together. A prosocial-choice task can fail because the actor does not value the recipient’s outcome, because food competition suppresses action, because the partner is socially risky, because the action–outcome mapping is too abstract, because recipient protest changes later social cost, or because the chosen option is biased by apparatus structure. Without separating these components, social modulation may be underestimated, overestimated, or attributed to the wrong mechanism.

The proposed framework addresses a central tension in the primate helping literature. On one hand, outcome labels such as altruism, empathy, or prosociality are overinterpreted when they are treated as mechanisms, a concern consistent with broader reviews distinguishing selfish, mutualistic, and other-regarding motivations in primate prosociality (de Waal and Suchak, 2010). On the other hand, this scepticism does not imply that primate helping studies lack scientific value. Their value lies in showing how different task designs isolate different components of social action selection. Instrumental-helping tasks often maximise visibility, goal clarity, and affordance clarity. Tool-transfer tasks are more diagnostic when they require selection of the appropriate object for the partner’s visible problem. Prosocial food-choice tasks introduce competition, abstract action–outcome mappings, social partner effects, recipient signalling, and motor or apparatus biases. Cooperative breeders may differ not because they possess a qualitatively different moral faculty, but because their socio-ecology tunes recipient-value weighting, relationship value, cost tolerance, tolerance of co-feeding, and competition differently. The framework therefore shifts the question away from whether a species is prosocial in general and toward which task, partner, temporal, and valuation conditions allow recipient benefit to influence action. This account also clarifies how stronger evidence would be obtained. A claim about other-regarding valuation should survive comparison against motor-bias, self-interest, cost, competition, partner-identity, and relationship models. A claim about empathy-like responding should require counterfactual sensitivity to inferred need after other variables are controlled. A claim about costly other-benefit should separate the cost structure of the outcome from the motivation that produced it, and a stricter altruism-like claim should require negative net actor outcome. A claim about collaboration should show non-additive joint payoff and sensitivity to partner action or reliability, whereas helping in collaborative and non-collaborative contexts should be separated analytically (Greenberg et al., 2010). A claim about socially mediated prosociality should show that partner behaviour changes subsequent choices in a way that cannot be reduced to side bias, action repetition, food motivation, or immediate solicitation alone.

The model is intentionally minimal. It does not solve the full problem of social cognition in nonhuman primates. It does, however, impose useful discipline. It replaces a vocabulary of psychologically ambiguous constructs with a common computational architecture in which those constructs become output labels, task presets, or parameter-dependent special cases. This is the main theoretical contribution: the same model can generate helping, prosociality, costly other-benefit, strict altruism-like outcomes, empathy-like responding, and collaboration without assuming that any of these labels is a basic psychological cause.

### Implications for comparative cognition

Although the present model is not a comparative species model, it is designed to make comparative hypotheses more precise. Rather than asking whether one species is more altruistic or prosocial than another, species or social systems can be compared in terms of parameter distributions: sensitivity to recipient benefit, cost tolerance, competition, relationship value, solicitation, affordance clarity, and motor or apparatus bias. This is also why social determinants such as partner identity, tolerance, affiliation, or dominance relation are not secondary complications. They define the conditions under which a detected recipient need becomes behaviourally relevant. Comparative cognition therefore enters not as a ranking of species, but as a framework for identifying which task and valuation components differ across taxa, dyads, or socio-ecological contexts.

### Limitations and next steps

The model is intended as a generative proof of principle rather than as a fitted empirical model. The parameter presets should therefore not be interpreted as species estimates. They describe regions of model space that can later be constrained by data. The current simulation also uses a simplified two-action structure, whereas real social tasks often contain multiple possible actions, repeated interactions, partner feedback, and changing motivational states. The literature coverage is selective rather than systematic. Its purpose is to identify recurring inferential problems and to show how they can be formalised. The next empirical step is to design tasks in which the gates are independently measured or manipulated. Need visibility, request intensity, affordance understanding, action cost, competition, relationship value, and motor bias should be estimated outside the focal helping outcome whenever possible. The computational framework can then be used for model comparison: other-outcome sensitivity should improve prediction beyond bias, self-interest, cost, competition, and relationship terms before stronger psychological interpretations are made.

## Conclusion

Helping behaviour should not be interpreted as a direct behavioural readout of altruism, empathy, concern, or other-regarding preference. It should be treated as the endpoint of socially gated action selection. Recipient need must be detected, the recipient’s goal must be inferred, an effective helping affordance must be recognised, and the expected value of action must overcome cost, competition, dominance risk, relationship constraints, and motor or apparatus bias. Traditional psychological labels are then assigned to the resulting action–outcome pattern. They should not be used as the starting explanation for that pattern. The central contribution is therefore inferential rather than motivational. The model does not solve the problem of helping motivation in nonhuman primates, nor does it identify altruism, empathy, or prosocial preference from behaviour alone. It constrains the kinds of claims that can be made from particular experimental designs. Stronger interpretations require diagnostic contrasts showing that behaviour tracks recipient need or recipient outcome after self-interest, cost, competition, relationship value, request context, partner feedback, and response bias have been separated. The claim is not that altruism, empathy, helping, prosociality, and collaboration are identical. They are not. The claim is that they can be generated as different outcome profiles of one socially gated action-selection architecture. For primate cognition, this shifts the target of explanation away from whether a species is prosocial in general and toward the task, partner, and valuation conditions under which recipient benefit is allowed to influence action.

## Acknowledgements

The author thanks colleagues and students whose discussions of primate helping, prosocial choice, and comparative cognition motivated parts of the framework.

## Statements and Declarations

### Ethics approval

No live animals, human participants, biological material, or privately held behavioural data were used. The study develops a theoretical model and analyses simulated data only. Under the applicable institutional rules for theoretical and simulation-based research at Taipei Medical University, no animal ethics approval was required.

### Code and data availability

The MATLAB code used to generate the simulations, figures, and output summaries is available in the public GitHub repository: https://github.com/ChristophDahl/social-gating-prosocial-behaviour.

### Author contributions

Christoph D. Dahl conceived the study, developed the model, implemented the simulations, interpreted the results, and wrote the manuscript.

