## Supplementary information for "Primate Prosocial Behaviour as Socially Gated Action Selection"

### Supplementary Material for:
#### Primate Prosocial Behaviour as Socially Gated Action Selection

Christoph D. Dahl
Graduate Institute of Mind, Brain and Consciousness, Taipei Medical University, Taipei, Taiwan

This supplementary material provides expanded literature and task-preset tables that support the main manuscript's treatment of prosocial behaviour as socially gated action selection without interrupting the central model–simulation–interpretation sequence. It also provides simulation-documentation tables and one supplementary failure-decomposition figure derived from the MATLAB implementation and output files.

### Supplementary Table S1. Empirical background and paradigms

Table S1 expands the literature mapping used to motivate the social-gating framework. The table is intended as an interpretive map rather than a comprehensive review.

**Table S1:** Empirical literatures reorganised as social-gating problems. Different prosocial and helping-related paradigms can be interpreted as tests of distinct gates or valuation terms rather than as direct assays of altruism, empathy, or prosocial motivation.

| Literature class | Representative paradigms | Main gate or model term | Main interpretive ambiguity |
|---|---|---|---|
| Instrumental helping | Reaching tasks, object access, removal of simple obstacles. | Need detection; goal transparency; affordance clarity; low action cost. | Helping may be frequent because the task is transparent and cheap, not because strong other-regarding motivation is required. |
| Tool transfer | Visible or hidden recipient problem; transfer of a useful object or tool. | Affordance mapping; visual access to need; solicitation sensitivity. | Targeted transfer is informative, but helping may depend strongly on request signals and visibility of the partner's problem. |
| Prosocial-choice tasks | Token choice, food-delivery apparatuses, touch-screen-guided choices, one-option versus two-option giving tasks. | Other-outcome valuation; partner identity; relationship value; competition; apparatus bias; motor lateralisation; side preference. | Apparent prosociality can vary with social context and partner-related variables (Mendonça et al., 2018), but can also be produced or distorted by token placement, reaching convenience, or response bias (Sikorska et al., 2026). |

*Continued on next page*

**Table S1:** Empirical literatures reorganised as social-gating problems (continued).

| Literature class | Representative paradigms | Main gate or model term | Main interpretive ambiguity |
|---|---|---|---|
| Food sharing and tolerated taking | Food transfer, co-feeding tolerance, tolerated theft, harassment-induced sharing. | Action cost; resource competition; relationship value; dominance risk; reciprocal history. | Recipient benefit does not identify prosocial motivation, because sharing can arise from tolerance, harassment avoidance, kinship, dominance, reciprocal history, or partner value (Stevens, 2004; Jaeggi and van Schaik, 2011; Jaeggi and Gurven, 2013). |
| Consolation and post-conflict affiliation | Third-party affiliation after aggression; contact directed to victims or distressed individuals. | Inferred social or affective state; relationship value; arousal regulation. | Post-conflict contact may reflect concern for another's state, but may also reflect self-arousal reduction, relationship repair, or tension regulation. |
| Cooperation and collaboration | Cooperative pulling, partner choice, role coordination, partner reliability tasks. | Joint payoff; partner prediction; action interdependence; tolerance constraints. | Successful coordination may reflect mutualism, direct self-benefit, partner choice, tolerance constraints, or sensitivity to partner competence rather than other-regarding motivation (Melis et al., 2006a,b; Greenberg et al., 2010). |
| Cooperative breeding and provisioning | Food provisioning in callitrichids; infant-care and group-care contexts. | Other-outcome weighting; low competition; care ecology; relationship value. | Stronger provisioning need not imply a special prosocial faculty; it may reflect socio-ecological tuning of cost, tolerance, care ecology, and recipient-value parameters (Burkart et al., 2007; Jaeggi et al., 2010). |

#### Supplementary Table S2. Task-preset contrasts

Table S2 expands the task-preset logic used in the illustrative simulations. It states the diagnostic contrast for each prosocial, helping-related, or collaborative task preset and the expected bottleneck under the social-gating account.

**Table S2:** Canonical task presets expressed as model contrasts. The table clarifies the diagnostic logic of each task preset rather than reproducing all numerical values used in the simulation. Full numerical task-preset and parameter-preset values are reported in Tables S4 and S6.

| Task preset | Main model contrast | Expected bottleneck | Diagnostic prediction |
|---|---|---|---|
| Instrumental reaching | Recipient need and helping affordance are both visible; action cost is low. | Valuation rather than competence. | Helping should be frequent when even moderate other-outcome sensitivity is present. Low helping would be informative only if need visibility, affordance clarity, cost, and action access are all controlled. |
| Visible tool transfer | Recipient need and the useful object or action are observable to the actor. | Cost, valuation, and request sensitivity. | Targeted helping should be high, especially when requests are present and the useful object is clearly linked to the recipient's problem. |
| Hidden tool transfer | Request may be present, but recipient need, goal, or the relevant apparatus state is partly occluded. | Need detection and affordance mapping. | Helping should decline relative to visible tool transfer despite similar action cost. A null result may reflect hidden need or unclear affordance rather than lack of other-regarding valuation. |
| Food token, randomised | Recipient benefit is abstract, but side and position bias are reduced by randomisation. | Competition, abstract action–outcome mapping, and weak other-outcome valuation. | Prosocial choice should be modest unless recipient-benefit valuation survives food competition and the actor understands the token–outcome relation. |
| Food token, fixed position | Recipient benefit is abstract and token location can covary with physical reaching convenience. | Bias, competition, affordance ambiguity, and motor lateralisation. | Apparent prosociality can be unstable or artefactual if token placement, hand preference, side preference, or reaching convenience is uncontrolled. |

*Continued on next page*

| Task preset | Main model contrast | Expected bottleneck | Diagnostic prediction |
|---|---|---|---|
| Apparatus bias, no benefit | The help-like option is strongly favoured by motor or apparatus bias, but recipient benefit is absent. | False-positive response tendency. | Help-like choice can be high while actual helping remains zero. This preset separates nominal prosocial choice from realised recipient benefit. |
| Low-competition provisioning | Recipient benefit is clear and competition is weak. | Other-outcome valuation and relationship tolerance. | Provisioning should increase when other-outcome weighting is high, competitive suppression is low, or tolerance reduces the effective cost of allowing recipient benefit. |
| High-cost help | Recipient benefit is clear, but action is costly to the actor. | Cost override. | Helping should occur only when other-outcome valuation, relationship value, reciprocity, or solicitation outweighs direct cost. Low helping does not by itself imply failure to understand the recipient's need. |
| Dominance-risk help | Recipient benefit is clear, but action carries social risk or may provoke competition. | Cost, dominance risk, and competition. | Helping should be strongly suppressed except when other-outcome valuation or relationship value is high enough to overcome social risk. |
| Reliable-partner collaboration | Actor and recipient both benefit from coordinated action. | Partner reliability and joint payoff. | Action should depend on expected joint payoff and partner reliability rather than recipient benefit alone. Partner-benefiting action is therefore not necessarily altruism-like. |

### Mathematical note on Equations 17 and 18

Equation 17 in the main manuscript expresses a partial derivative. A partial derivative asks how one quantity changes when one input is varied while other relevant inputs are held fixed. Here, the quantity of interest is the probability that the actor selects the recipient-benefiting action, $P_i(a_{\text{help}} \mid z_t)$. The varied input is inferred recipient need, $\hat{N}_{ij,t}$. Thus,

$$\frac{\partial P_i(a_{\text{help}} \mid z_t)}{\partial \hat{N}_{ij,t}} > 0$$

means that increasing inferred recipient need should increase the probability of choosing the recipient-benefiting action, after direct actor benefit, action cost, competition, relationship value, solicitation, and motor or apparatus bias have been held constant. In the two-action simulation, this can be seen directly from the softmax rule. If the actor chooses between a recipient-benefiting action $a_{\text{help}}$ and a non-recipient-benefiting action $a_0$, then

$$P_i(a_{\text{help}}) = \frac{\exp(\beta U_i(a_{\text{help}}))}{\exp(\beta U_i(a_{\text{help}})) + \exp(\beta U_i(a_0))}.$$

This can be rewritten as

$$P_i(a_{\text{help}}) = \frac{1}{1 + \exp[-\beta (U_i(a_{\text{help}}) - U_i(a_0))]}.$$

Thus, the probability of helping increases when the utility difference $U_i(a_{\text{help}}) - U_i(a_0)$ increases. In the model, inferred need enters the utility of the recipient-benefiting action through the social gate,

$$\Gamma_{ij,t}(a) = \hat{N}_{ij,t} Q_{ij,t}(a).$$

For a recipient-benefiting action, the part of the utility that depends on inferred need is therefore

$$\hat{N}_{ij,t} Q_{ij,t}(a) \left(w_{\text{other}} + w_{\text{rel}} Rel_{ij} + w_{\text{recip}} Recip_{ij} + w_{\text{sol}} S_t\right) \widehat{\Delta X}_{j,t}(a).$$

The derivative of helping probability with respect to inferred need is therefore proportional to

$$Q_{ij,t}(a) \left(w_{\text{other}} + w_{\text{rel}} Rel_{ij} + w_{\text{recip}} Recip_{ij} + w_{\text{sol}} S_t\right) \widehat{\Delta X}_{j,t}(a).$$

More explicitly, in the two-action case,

$$\begin{aligned} \frac{\partial P_i(a_{\text{help}})}{\partial \hat{N}_{ij,t}} &= \beta P_i(a_{\text{help}}) \left[1 - P_i(a_{\text{help}})\right] Q_{ij,t}(a_{\text{help}}) \\ &\quad \times \left(w_{\text{other}} + w_{\text{rel}} Rel_{ij} + w_{\text{recip}} Recip_{ij} + w_{\text{sol}} S_t\right) \widehat{\Delta X}_{j,t}(a_{\text{help}}). \end{aligned}$$

This expression clarifies the diagnostic meaning of Equation 17. Need sensitivity is expected only when an effective helping affordance is represented, when the recipient-benefit term has positive weight, and when the action is expected to improve the recipient's state. A recipient-benefiting outcome alone is therefore insufficient. The model asks whether the probability of that action changes when inferred need is counterfactually varied.

Equation 18 expresses a different mathematical idea: an interaction between the actor's action and the partner's action. In continuous notation, this is written as a mixed partial derivative,

$$\frac{\partial^2 X_{ij}}{\partial a_i \partial a_j} \neq 0.$$

The expression asks whether the effect of the actor's action depends on the partner's action. If the actor's action has the same value regardless of what the partner does, then there is no interaction and the mixed derivative is zero. If the value of the actor's action changes depending on the partner's action, then the mixed derivative is nonzero. Because most behavioural tasks involve discrete actions rather than infinitesimal changes in action, the same idea can be written as a finite-difference interaction contrast. Suppose each individual can either not act, coded as 0, or act, coded as 1. The joint outcome can then be written as $X_{ij}(a_i,a_j)$. The interaction contrast is

$$\Delta^2 X_{ij} = X_{ij}(1,1) - X_{ij}(1,0) - X_{ij}(0,1) + X_{ij}(0,0).$$

If

$$\Delta^2 X_{ij} = 0,$$

then the actor's and partner's contributions are additive. The joint outcome can be decomposed into separate individual effects, and no special joint-payoff term is required. If

$$\Delta^2 X_{ij} \neq 0,$$

then the outcome contains a non-additive joint component. This is the formal signature of collaboration in the present framework: the value of one individual's action depends on the coordinated action of the other individual. Equation 18 is therefore not intended to imply that animal actions vary continuously. It is a compact notation for the same interaction logic captured by the finite-difference contrast above.

### Supplementary simulation documentation

The following tables document the simulation fields, preset values, and generated output files. Table S3 defines the task-preset variables used to construct the simulated task presets. Table S4 reports the full numerical task-preset matrix used in the simulation. Table S5 defines the parameter-preset fields. Table S6 reports the full numerical parameter-preset matrix. Table S7 specifies the output-summary columns generated from `summary_by_task_and_system.csv`.

**Table S3:** Task-preset fields used to define each simulated task preset.

| Preset field | Interpretation |
|---|---|
| `task` | Name of the task preset used for this simulation run. |
| `visibilityNeed` | Visibility of the recipient's need. |
| `solicitation` | Request or solicitation intensity. |
| `goalTransparency` | Transparency of the recipient's goal. |
| `affordanceClarity` | Clarity of the helping action–outcome mapping. |
| `familiarity` | Actor's familiarity with the task or apparatus. |
| `occlusion` | Occlusion or ambiguity of the recipient's problem or apparatus state. |
| `actionCost` | Cost of the helping action to the actor. |
| `competition` | Resource conflict, dominance risk, or social competition. |
| `relationshipValue` | Relationship, tolerance, kinship, affiliation, or partner-value term. |
| `reciprocalHistory` | Reciprocal or interaction-history term. |
| `motorBias` | Motor, side, hand, token-position, or apparatus bias favouring the help-like action. |
| `expectedSelfBenefit` | Direct actor benefit expected from the helping action. |
| `expectedOtherBenefit` | Expected improvement in recipient state. |
| `expectedJointBenefit` | Non-additive joint benefit, relevant for collaboration. |
| `actualOtherBenefit` | Realised recipient benefit if the help-like action is selected. |
| `actualSelfBenefit` | Realised direct actor benefit before subtracting action cost. |
| `isCollaborative` | Indicator for collaborative task structure with actor benefit, recipient benefit, and joint payoff. |

**Table S4:** Full task-preset matrix used in the illustrative simulation. Columns correspond to the variables defined in the model. The distinction between expected and actual self- and other-benefit prevents low-cost recipient-benefiting actions from being treated as self-benefiting unless a direct actor benefit is explicitly specified.

| Task preset | $V$ | $S$ | $G$ | $A$ | $F$ | $O$ | $C$ | $Comp$ | $Rel$ | $Recip$ | $E[\Delta X_i]$ | $E[\Delta X_j]$ | $J$ | $M$ | $\Delta X_j$ | $\Delta X_i$ | Collab. |
|---|---|---|---|---|---|---|---|---|---|---|---|---|---|---|---|---|---|
| Instrumental reaching | 0.95 | 0.70 | 0.90 | 0.92 | 0.82 | 0.05 | 0.10 | 0.05 | 0.35 | 0.10 | 0.00 | 0.90 | 0.00 | 0.00 | 0.90 | 0.00 | 0 |
| Visible tool transfer | 0.96 | 0.75 | 0.90 | 0.90 | 0.78 | 0.04 | 0.14 | 0.05 | 0.35 | 0.12 | 0.00 | 0.88 | 0.00 | 0.00 | 0.88 | 0.00 | 0 |
| Hidden tool transfer | 0.25 | 0.65 | 0.45 | 0.45 | 0.58 | 0.75 | 0.14 | 0.05 | 0.35 | 0.12 | 0.00 | 0.82 | 0.00 | 0.00 | 0.82 | 0.00 | 0 |
| Food-token randomised | 0.45 | 0.18 | 0.42 | 0.55 | 0.55 | 0.25 | 0.05 | 0.55 | 0.15 | 0.05 | 0.00 | 0.65 | 0.00 | 0.00 | 0.65 | 0.00 | 0 |
| Food-token fixed position | 0.45 | 0.18 | 0.42 | 0.50 | 0.55 | 0.25 | 0.05 | 0.55 | 0.15 | 0.05 | 0.00 | 0.65 | 0.00 | -0.20 | 0.65 | 0.00 | 0 |
| Apparatus bias, no benefit | 0.35 | 0.00 | 0.25 | 0.35 | 0.55 | 0.35 | 0.02 | 0.00 | 0.00 | 0.00 | 0.00 | 0.00 | 0.00 | 2.25 | 0.00 | 0.00 | 0 |
| Low-competition provisioning | 0.66 | 0.20 | 0.62 | 0.65 | 0.70 | 0.12 | 0.08 | 0.10 | 0.45 | 0.15 | 0.00 | 0.75 | 0.00 | 0.00 | 0.75 | 0.00 | 0 |
| High-cost help | 0.96 | 0.55 | 0.90 | 0.92 | 0.82 | 0.04 | 0.80 | 0.15 | 0.35 | 0.10 | 0.00 | 1.00 | 0.00 | 0.00 | 1.00 | 0.00 | 0 |
| Dominance-risk help | 0.86 | 0.60 | 0.85 | 0.82 | 0.76 | 0.08 | 0.55 | 0.75 | 0.25 | 0.10 | 0.00 | 0.92 | 0.00 | 0.00 | 0.92 | 0.00 | 0 |
| Reliable-partner collaboration | 0.86 | 0.40 | 0.80 | 0.85 | 0.80 | 0.07 | 0.25 | 0.10 | 0.50 | 0.20 | 0.75 | 0.75 | 0.80 | 0.00 | 0.75 | 0.75 | 1 |

**Table S5:** Parameter-preset fields used to define each simulated parameter preset.

| Parameter field | Interpretation |
|---|---|
| `socialSystem` | Name of the parameter preset used for this simulation run. |
| $w_{self}$ | Weight on direct actor benefit. |
| $w_{cost}$ | Weight on action cost. |
| $w_{comp}$ | Weight on competition or resource conflict. |
| $w_{other}$ | Weight on socially gated recipient benefit. |
| $w_{rel}$ | Relationship-value weight. |
| $w_{recip}$ | Reciprocity or interaction-history weight. |
| $w_{sol}$ | Solicitation or request-sensitivity weight. |
| $w_{joint}$ | Weight on non-additive joint payoff. |
| $\beta$ | Softmax choice consistency. |
| `helpActionThreshold` | Action-initiation threshold for recipient-only helping actions. |
| `actorBenefitActionThreshold` | Reduced action-initiation threshold when direct actor benefit is present. |
| `actionBenefitTolerance` | Numerical tolerance used to distinguish direct actor benefit from recipient-only benefit. |

**Table S6:** Full parameter-preset matrix used in the illustrative simulation. The presets define model-space configurations rather than species models.

| Parameter preset | $w_{self}$ | $w_{other}$ | $w_{joint}$ | $w_{cost}$ | $w_{comp}$ | $w_{rel}$ | $w_{recip}$ | $w_{sol}$ | $\beta$ |
|---|---|---|---|---|---|---|---|---|---|
| Moderate other-weight / high competition sensitivity | 1.00 | 0.28 | 0.60 | 1.35 | 1.05 | 0.45 | 0.30 | 0.35 | 3.00 |
| High other-weight / moderate competition sensitivity | 1.00 | 0.95 | 0.70 | 1.00 | 0.55 | 0.65 | 0.35 | 0.45 | 3.20 |
| High other-weight / low competition sensitivity | 1.00 | 0.65 | 0.60 | 1.15 | 0.45 | 0.55 | 0.30 | 0.35 | 3.00 |
| Low other-weight | 1.00 | 0.05 | 0.25 | 1.60 | 1.10 | 0.10 | 0.10 | 0.10 | 3.00 |
| Pure self-interest | 1.00 | 0.00 | 0.00 | 1.80 | 1.20 | 0.00 | 0.00 | 0.00 | 4.00 |

**Table S7:** Columns in the task-by-parameter output summary generated from `summary_by_task_and_system.csv`.

| Output field | Interpretation |
|---|---|
| `task`, `socialSystem`, `n` | Task preset, parameter preset, and number of simulated trials. |
| `choiceHelpRate` | Proportion of trials in which the help-like action was selected. |
| `helpingRate` | Proportion of trials in which the selected action improved the recipient's state. |
| `prosocialRate` | Proportion of low-cost recipient-benefit outcomes. |
| `costlyOtherBenefitRate` | Proportion of recipient-benefit outcomes with actor cost. |
| `strictAltruismLikeRate` | Proportion of recipient-benefit outcomes with negative net actor outcome. |
| `collaborationLikeRate` | Proportion of collaboration-like outcomes in the joint-payoff preset. |
| `falsePositiveRate` or `falsePositiveBiasRate` | Proportion of help-like choices without recipient benefit. |
| `meanPNeed`, `meanPAfford`, `meanGate`, `meanPHelp` | Mean inferred need, affordance recognition, social gate, and helping-action probability. |
| `needFailureRate`, `affordanceFailureRate` | Failure rates at the need-inference and affordance-recognition stages. |
| `costCompetitionOverrideRate` | Failure rate due to cost, risk, or competition override. |
| `valuationNoHelpRate` | Failure rate due to valuation-based non-helping. |
| `successRate` | Proportion of trials reaching the realised helping outcome. |

### Additional simulation checks

**Model-ablation scores.** Model-ablation scoring was used as a component-ablation check rather than as independent empirical model validation. The purpose was to ask whether simpler candidate accounts could reproduce the choices generated by the full social-gating architecture. Bias-only, self-interest, competition-only, and relationship-only variants were therefore compared with the full model using the generated choice data. The expected pattern was recovered: when other-outcome sensitivity was present, removing social-gating components reduced predictive adequacy, whereas in the pure self-interest control simpler non-social variants were not penalised in the same way. This confirms that the ablation analysis behaves as an internal diagnostic of the simulated architecture, not as an empirical test of whether the architecture is true of animal behaviour.

**False-positive apparatus-bias check.** The apparatus-bias/no-benefit preset was included to demonstrate a pure false-positive condition. In this preset, the help-like option was strongly favoured by motor or apparatus bias, but selecting it did not improve the recipient's state. Nominal help-like choice was therefore high across parameter presets, whereas realised recipient benefit, prosocial choice, costly recipient benefit, and strict altruism-like outcomes remained zero. This check illustrates why the selected option in a prosocial-choice task should not be interpreted as recipient benefit unless side, hand, location, reaching convenience, and actual recipient outcome are separated.

### Supplementary figure

Figure S1 expands the main manuscript's averaged failure-decomposition figure by showing the same decomposition separately for each parameter preset.

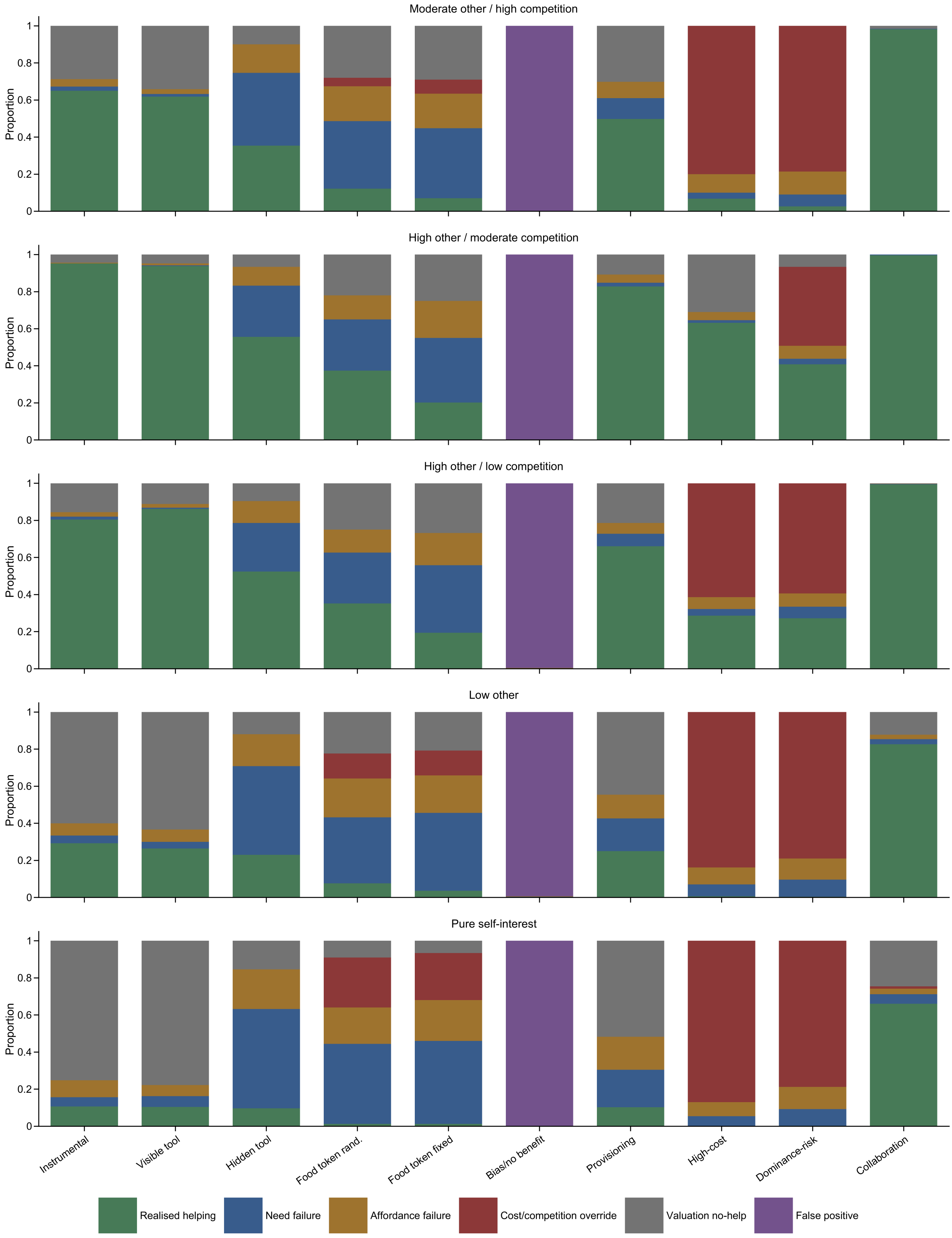


**Figure S1:** Failure decomposition by task preset and parameter preset. The plot expands the main manuscript's averaged failure-decomposition figure by showing how need-inference failure, affordance failure, cost/competition override, valuation-based non-action, false-positive bias, and realised recipient benefit vary across parameter presets.
